# Mathematical Foundations of Beta Diversity: Why Common Metrics Fail in Microbiome Analysis

**DOI:** 10.1101/2025.08.14.670326

**Authors:** Zihan Zhu, Yiqian Zhang, Wenhao Li, Michael Greenacre, Satabdi Saha, Yushu Shi, Liangliang Zhang

## Abstract

**Background:** In microbiome studies, beta diversity quantifies dissimilarity between samples and is often visualized using ordination techniques. It enables researchers to characterize ecological structure, compare microbial communities, assess environmental or host-driven heterogeneity, and track longitudinal shifts over time. Although many diversity indices were originally developed with practical goals in mind, they lack a unified framework to ensure theoretical rigor and validity. This gap makes it challenging for researchers to evaluate and select appropriate beta diversity measures for microbiome analyses, potentially leading to biased analyses and invalid conclusions.

**Results:** To bridge the persistent knowledge gaps, we systematically evaluate the commonly used beta diversity measures according to key mathematical properties, including whether they are true metrics, conform to Euclidean geometry, and satisfy conditional negative definiteness. We show that their violations can compromise downstream analyses such as PCoA, PERMANOVA, and kernel-based tests. In addition, drawing on mathematical consensus, we introduce a novel four-category classification of beta diversity measures: scale difference, difference scale, Hamming difference, and distribution difference. Complementing this framework, we build diagnostic tools for assessing Euclidean validity and develop remedial strategies that correct problematic dissimilarity matrices while preserving ordination structures. We demonstrate the effectiveness of these solutions using real-world microbiome datasets.

**Conclusions:** These results establish a unified framework for evaluating beta diversity in microbiome research, supported by an R package, interactive Shiny app, and step-by-step tutorials. The framework provides a clear roadmap for selecting and refining dissimilarity metrics, paving the way for future methodological advances.

## 1 Introduction

Microbiome research differs fundamentally from traditional human omics studies, which focus on molecular data (e.g. genomics, transcriptomics, or proteomics) derived from a single organism. In contrast, microbiome studies investigate a complex, multi-species community of microorganisms inhabiting diverse environments. These communities exhibit intricate ecological interactions—such as mutualism, competition, synergism, and antagonism—that influence both the microbial composition and host physiology [1]. To quantitatively capture the structure and variability of these communities, researchers rely on diversity indices rooted in ecological theory [2], while developing new metrics tailored to the unique features of microbiome data, such as phylogenetic relatedness and compositionality. As such, ecological principles serve as a foundational lens for understanding microbial ecosystems [3].

Among these ecological principles, understanding how biological communities vary across space and time has long been a central theme in ecology. Whittaker’s alpha–beta–gamma framework [4] formalized this idea by partitioning biodiversity within and between habitats, establishing the foundation for modern diversity metrics. In particular, beta diversity captures the turnover or differentiation among communities across varying environments or conditions [5, 6]. In the context of microbiome research, it provides a fundamental means to assess how microbial community structures differ across individuals, body sites, or environmental conditions, thereby linking ecological variability to underlying biological or environmental drivers.

Although many dissimilarity measures were developed for practical purposes and have proven invaluable in ecological and biomedical research, problems arise when they fail to satisfy key mathematical properties. In such cases, the resulting dissimilarities may fail to represent the true spatial relationships between samples. This non-metric nature can introduce distortions to downstream analyses such as ordination, clustering, or kernel-based models, ultimately affecting the interpretation of community dissimilarity patterns. To illustrate this issue, we present a simple example highlighting the impact of non-metric dissimilarities on principal coordinates analysis (PCoA) visualizations. As shown in Figure 1, violations of metric properties can distort spatial relationships between samples, making dissimilar communities appear artificially close in reduced-dimensional space. We refer to this phenomenon as the illusion of similarity (IOS), which quantifies mismatches between true and projected neighborhood structures. In this toy example, the average IOS score reaches as much as 0.7, highlighting the severity of geometric distortion that can arise when non-Euclidean dissimilarities are used for ordination or clustering.

**Fig. 1:**
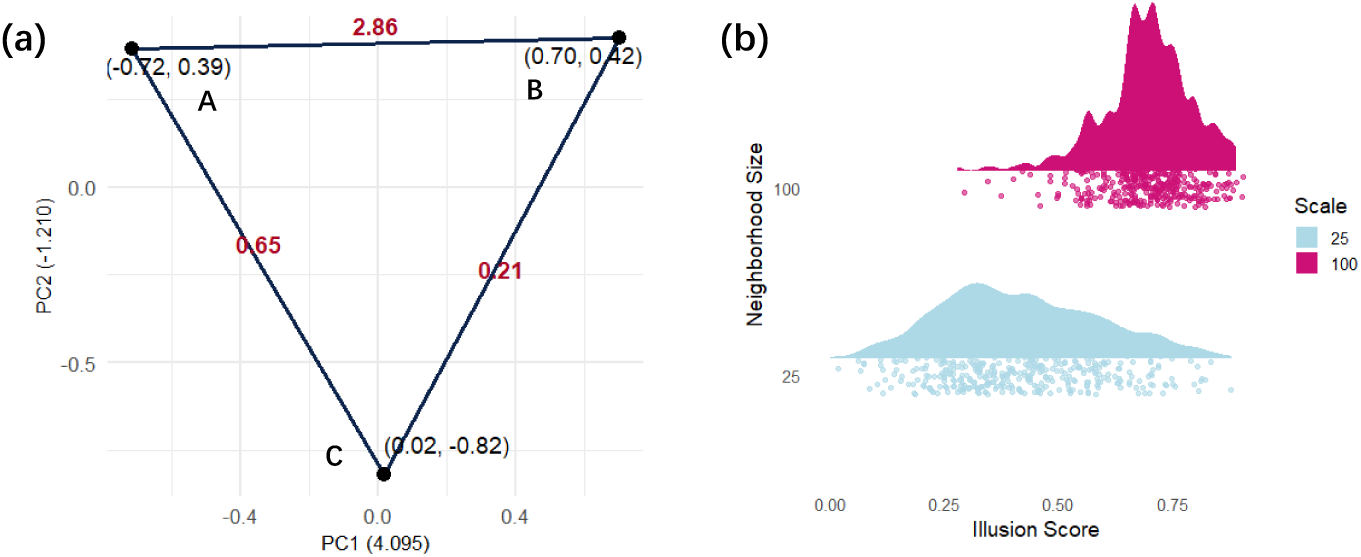
Illustration of geometric and statistical distortions due to non-Euclidean dissimilarities based on a Bray–Curtis dissimilarity matrix among samples from a problematic dataset (to be introduced in Section 4.1). (a) A toy example illustrating the distortion and discrepancy between the reduced space and the original distances caused by the non-metric property. Pairwise Bray–Curtis dissimilarities were calculated and are shown as red numbers between samples. The three pairwise distances violate the triangle inequality, illustrating the non-metric property of the measure. Principal Coordinates Analysis (PCoA) was then performed on the dissimilarity matrix, yielding a negative second eigenvalue. The coordinates of each sample are labeled at the vertices, showing that distances appear equal in the reduced space, although they are not in the original dissimilarity space. (b) Raincloud plot showing the distribution of the illusion of similarity (IOS) score across neighborhood sizes. IOS quantifies the fraction of non-true neighbors introduced by ordination; the mean IOS value (≈ 0.69) reflects the magnitude of geometric distortion caused by non-metric dissimilarities.

The above example illustrates a broad and serious issue in microbiome research. Despite growing use of beta diversity in microbiome studies [7], many studies lack a unified framework to assess the theoretical validity of these measures. Without a rigorous foundation and evaluation framework, it becomes difficult for researchers to assess or select appropriate beta diversity metrics—raising the risk of analytical distortions, biased results, and misleading biological conclusions. To bridge these persistent knowledge gaps, we systematically review and evaluate the commonly used beta diversity measures according to key mathematical properties, including whether they are true metrics, conform to Euclidean geometry, and satisfy conditional negative definiteness. We show that their violations can compromise downstream analyses such as PCoA, PERMANOVA [8], and kernel-based methods. To guide robust and interpretable analyses, we introduce a unified classification framework, develop diagnostic tools to assess distortion, and provide practical recommendations for selecting or adjusting dissimilarity measures. We demonstrate the effectiveness of these solutions using real-world microbiome datasets.The resulting framework is implemented in an R package with an interactive Shiny application and step-by-step tutorials to support broad adoption. The remainder of this paper is organized as follows. Section 2 reviews the theoretical background and recent developments of beta-diversity measures. Section 3 introduces three key mathematical properties that ideal dissimilarities should satisfy and presents a classification framework that groups existing measures according to their shared theoretical features. Section 4 illustrates the proposed framework and diagnostic tools using multiple real-world microbiome datasets. Section 5 provides a general discussion and outlines directions for future research, and Section 6 concludes the paper.

## 2 Background

The origins of ecology as a science trace back to the application of experimental and mathematical approaches to understanding organism–environment interactions, community structure and succession, and population dynamics [9]. While early work focused on plants and animals, the development of high-throughput sequencing transformed our understanding of microorganisms, enabling comprehensive profiling of microbial communities [10, 11]. Numerous beta diversity measures have been proposed to characterize community turnover, each reflecting different assumptions about presence–absence, abundance, or phylogeny [12]. Classical indices such as Jaccard [13] and Sørensen [14] capture compositional differences, whereas modern metrics—such as UniFrac [15]—incorporate evolutionary relatedness to better reflect microbial lineage structure. These measures are now standard tools in ecological and microbiome data analysis, implemented in widely used software such as vegan and phyloseq.

A simple bibliometric study further demonstrates this trend: the proportion of microbiome studies with “beta diversity” mentioned in their title, abstract, or keywords has steadily increased over the past two decades (Supplementary Material Section 1). This growing prominence reflects the expanding scope of beta diversity analysis across microbiome applications. Plantinga and Wu [16] synthesized statistical frameworks for distance-based microbiome analysis, linking ecological theory with regression and hypothesis testing. Pasqualini et al. [17] demonstrated how statistical-physics models of beta diversity reveal stable and transitional gut microbiome states, while Simon et al. [18] illustrated its role in designing microbiome-based interventions for food safety and environmental health. Together, these studies underscore beta diversity as a bridge connecting ecological dynamics, host–microbe interactions, and applied microbiome science.

Despite its broad use, microbiome data introduce distinctive analytical challenges—compositional constraints, sparsity, and high dimensionality—that complicate distance estimation and inference [19]. Similar methodological concerns have been explored in other statistical contexts, such as approximation under limited replication [20], modeling incomplete or censored information [21], and scalable inference for complex dependencies [22]. The issue of selecting appropriate analytical models—highlighted by Meyners and Hasted [23] in sensory data—also parallels the need to align beta diversity metrics with ecological intent. In microbiome-specific research, Zhang et al. [24] proposed a unified framework combining proportion conversion and contrast transformation, offering a principled way to address compositionality and scaling effects in microbial abundance data. These developments motivate a systematic re-examination of the mathematical validity, geometric consistency, and interpretability of beta diversity measures, which is the focus of this study.

## 3 Method

We begin by introducing the concept of dissimilarity metrics, which serve as the foundation for quantifying beta diversity in microbial community analysis. Let *d*(*i, j*) denote a generic dissimilarity between microbial communities *i* and *j*, computed from input profiles such as species abundance, relative abundance, or presence–absence data. These pairwise values form the entries of a dissimilarity matrix *D* ∈ R*^N^*^×*N*^ , where each element satisfies

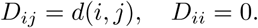

Such matrices serve as inputs for a variety of downstream analyses, including ordination techniques (e.g., PCoA), statistical tests (e.g., PERMANOVA [8], MiRKAT [25]), and community structure visualizations, which are used to reveal underlying ecological gradients.

In what follows, we detail the analytical principles and methodological choices that inform our evaluation and application of beta diversity measures. We first survey commonly used dissimilarities and their mathematical formulations. We then present a theoretical framework for evaluating key geometric and statistical properties—such as the metric condition, Euclidean embeddability, and conditional negative definiteness—which underpin many ecological interpretations. Next, we assess how deviations from these properties impact downstream analyses, and introduce diagnostic and correction tools to mitigate such effects. Together, these methods provide a comprehensive and geometry-aware foundation for evaluating beta diversity in microbiome studies.

### 3.1 An Overview of Commonly Used Dissimilarity Measures

Let’s first give a clear distinction: dissimilarity measures are like rulers—they provide the mathematical tools to measure the distances between microbial communities. Beta diversity, in contrast, is the landscape these rulers help us explore: it captures how community composition varies across environments or samples. For clarity and consistency in presenting mathematical formulas for each dissimilarity measure, we first define the notation and symbols used throughout. Let *N* denote the number of samples and *K* the number of observed taxa. For sample *i*, let *a_ik_* represent the absolute abundance (sequence read counts) of taxon *k*, with corresponding relative abundance 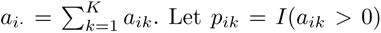 denote presence–absence indicators. As follows, we present dissimilarity measures in roughly chronological order and categorize them into two groups based on disciplinary origin: those developed primarily by biological researchers and those arising from quantitative fields such as mathematics and statistics. We begin with the biologically motivated measures. Additionally, following both prior literature and our own validation, we use the term distance exclusively for measures that satisfy all metric properties. All others are referred to as dissimilarities, irrespective of their original designation.

Table 1 provides an overview of representative beta diversity measures widely used in ecology and microbiome analysis. These metrics differ in their treatment of presence–absence, abundance, phylogenetic, and compositional information, as well as in their underlying geometric properties. Detailed derivations, historical contexts, applications, and full mathematical expressions for each measure are provided in Supplementary Material Section 2. The table summarizes the definitions that form the basis for Sections 3.2–3.4, where we build upon these concepts to conduct the systematic justification and construct the theoretical framework.

**Table 1:**
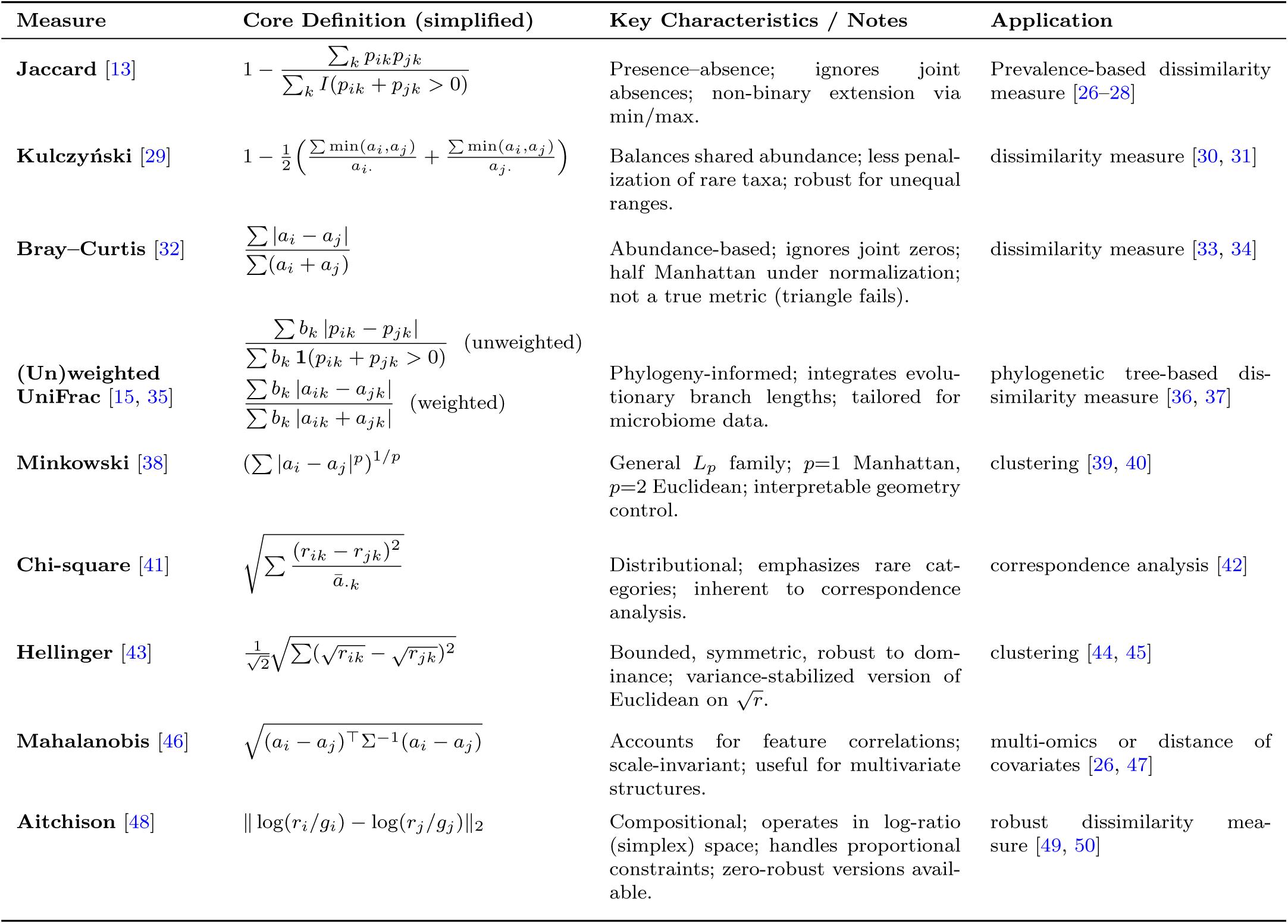
Summary of commonly used beta diversity measures in ecology and microbiome research.

### 3.2 Theoretical Properties of an Ideal Dissimilarity Matrix

To provide a mathematical justification and theoretical framework for existing beta diversity indices introduced above, we outline three fundamental mathematical properties that are often desired for an ideal dissimilarity matrix in microbial community analysis: the metric property, the Euclidean property, and conditional negative definiteness (CND). These properties form a hierarchy in terms of restrictiveness and analytical utility. While many indices satisfy the basic conditions of a metric, fewer are Euclidean or conditionally negative definite. Although the Euclidean property and CND are mathematically equivalent when applied to squared dissimilarities, they emphasize different perspectives: the Euclidean property supports geometric interpretations and ordination techniques such as principal coordinates analysis, while CND enables the construction of valid kernels for statistical learning and regression in Hilbert spaces. Discussing all three properties clarifies the theoretical capabilities and limitations of different dissimilarity measures and informs their appropriate use in ecological inference and downstream analysis.

#### 3.2.1 Metric Property

A dissimilarity matrix *D* ∈ R*^N^*^×*N*^ satisfies the metric property if its entries *D_ij_* fulfill the following four conditions for all *i, j, k* ∈ {1*, . . . , N* }:

1. Non-negativity:

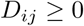

2. Identity of indiscernibles (reflexivity):

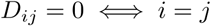

3. Symmetry:

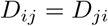

4. Triangle inequality:

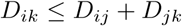

This set of conditions follows the definition of a metric in the sense of Gower [51]. These properties ensure that the dissimilarity measure functions in accordance with the principles of a mathematical distance. In microbial ecology, this means the index adheres to intuitive criteria such as identity (zero dissimilarity for identical communities), symmetry between community comparisons, and consistency when evaluating direct versus indirect differences. However, some widely used indices, such as Bray–Curtis dissimilarity, do not satisfy all four properties. In particular, they may violate the triangle inequality for certain sample pairs, and are therefore considered semi-metrics [52]. Although semi-metric dissimilarities remain meaningful and interpretable, their failure to satisfy the triangle inequality can limit their applicability in distance-based statistical methods that assume metric properties, such as metric multidimensional scaling (mMDS) [53].

Ecological interpretations of the triangle inequality arise naturally from fundamental principles of community assembly. A central hypothesis is the individualistic hypothesis, originally proposed by Whittaker [54], which posits that species respond independently to underlying environmental gradients, such as changes in soil pH (e.g., from 5 to 8), altitude (e.g., from 1000 to 3000 meters), or host diet (e.g., from high-fiber to high-fat intake). This hypothesis implies that changes in community composition occur gradually along such gradients, forming ecological continua. In this context, beta diversity—as a measure of dissimilarity between communities—should reflect this gradualism: if community B lies between A and C along a gradient, its compositional dissimilarity to both A and C should be more than A and C are to each other. This logic implies that the dissimilarity between A and C is bounded by the sum of dissimilarities involving B—a property exactly formalized by the triangle inequality. Further support comes from the idea of ecological continuity, especially in long-standing, undisturbed habitats. Odum [55] suggested that such continuity promotes gradual shifts in species composition over time or space, reinforcing the notion that community transitions are typically smooth rather than abrupt breaks or gaps in between. Thus, the triangle inequality aligns not merely with a mathematical structure, but with ecological expectations about how communities respond to continuous environmental variation. In microbial ecology, this idea translates into hypotheses about how microbiomes vary across gradients of host phenotype, tissue niche, or environmental exposure. For example, if a set of gut microbiomes vary along a diet-based continuum, then samples from intermediate diets should bridge the dissimilarities between more extreme ones. Consequently, observing violations of the triangle inequality in beta diversity matrices may signal ecological discontinuity, hidden host stratification, or methodological artifacts, making it an important diagnostic tool for both theoretical and practical purposes.

#### 3.2.2 Euclidean Property

A dissimilarity matrix *D* is said to have the Euclidean property if there exists an embedding of the *N* communities into a Euclidean space R*^m^*, for some 1 ≤ *m* ≤ *N* − 1, such that the squared dissimilarities correspond exactly to squared Euclidean distances between embedded points. That is, there exists a set of points {*x*_1_*, x*_2_*, . . . , x_N_* } ⊂ R*^m^*such that

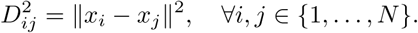

The concept of Euclidean embedding has deep mathematical roots. In 1841, Cayley introduced the Cayley–Menger determinant, providing early criteria for embedding point configurations in Euclidean space. Menger later formalized the necessary and sufficient conditions for a finite metric space to admit such an embedding [56]. A dissimilarity matrix that fulfills this criterion is said to have the Euclidean property, meaning that its pairwise values can be faithfully represented as distances in a Euclidean space without geometric distortion.

This property is particularly crucial for ordination methods in beta diversity analysis. When a dissimilarity matrix is Euclidean, techniques such as Principal Coordinates Analysis (PCoA) can accurately map the samples into a low-dimensional Euclidean space, preserving the dissimilarity structure as faithfully as possible. In this setting, the visualized distances between points directly correspond to the original dissimilarities. This allows researchers to intuitively interpret ecological gradients and clustering patterns in the data. In contrast, if the matrix lacks the Euclidean property, the embedding may introduce distortions, leading to misleading visual representations of community relationships [51, 57]. Thus, the Euclidean property is essential not only for mathematical consistency but also for ensuring meaningful ecological interpretation in ordination and visualization.

#### 3.2.3 Conditional Negative Definiteness

In mathematics, a symmetric function^1^ *f* : X × X → R is said to be conditionally negative definite (CND), if for any finite set of points {*x*_1_*, x*_2_*, . . . , x_N_* } ⊆ X and any real coefficients {*c*_1_*, . . . , c_N_* } satisfying 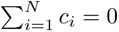, the following inequality holds:

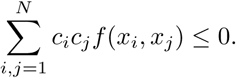

In beta diversity analysis, we often consider a squared dissimilarity matrix *D*^(2)^ = [*D*^2^ ]. It is said to be conditionally negative definite if

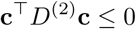

for all real vectors **c** = (*c*_1_*, . . . , c_N_* )^⊤^ with 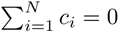.

The CND property plays a central role in ecological data analysis, particularly due to its connection with ordination. Many widely used dissimilarity measures, such as Rao’s quadratic entropy [58] and variance-based dissimilarities [59], are derived from conditionally negative definite functions. When the squared dissimilarities satisfy the CND condition, the corresponding matrix can be embedded into a Euclidean space without introducing curvature distortions.

Several ordination techniques, including Principal Coordinates Analysis (PCoA) and Double Principal Coordinates Analysis (DPCoA) [60], explicitly require this property. Notably, even when the original dissimilarity matrix *D* is not CND, its squared version *D*^(2)^ may be—this holds, for example, the Manhattan distance. This distinction is particularly relevant when evaluating whether a dissimilarity measure supports faithful Euclidean embedding. We further explore this connection in the next subsection.

#### 3.2.4 Relationship among Metric, Euclidean and CND Properties

The relationship among the metric property, the Euclidean property, and conditional negative definiteness (CND) has been studied extensively, beginning with Menger’s characterization theorem [61]. This theorem provides necessary and sufficient conditions for a semi-metric space to be isometrically embeddable in a Euclidean space, based on the positive semi-definiteness (PSD) of the associated squared distance matrix. This result was later generalized by Gower [51], who formalized the equivalence between the Euclidean property and the CND property for dissimilarities. Specifically, a dissimilarity measure is Euclidean if and only if its squared dissimilarity matrix is conditionally negative definite. Thus, the Euclidean and CND properties are mathematically equivalent. Furthermore, Blumenthal [62, 63] proved that if a semi-metric satisfies the Euclidean property, it must also satisfy the triangle inequality, and therefore is a metric. As most dissimilarity measures are at least semi-metrics, this result implies that every Euclidean dissimilarity is also a metric. However, the converse does not hold: not all metric dissimilarities are Euclidean. That is, some metric spaces cannot be isometrically embedded into any Euclidean space [64].

These theoretical relationships allow us to categorize dissimilarity measures into three levels of increasing mathematical adequacy:

1. Semi-metric dissimilarity: satisfies non-negativity, reflexivity, and symmetry, but may violate the triangle inequality.
2. Non-Euclidean metric dissimilarity: satisfies all metric properties, but not Euclidean.
3. Euclidean dissimilarity: both metric and Euclidean (i.e., squared dissimilarities are CND).

In practice, dissimilarity measures that are both metric and Euclidean are generally preferred, as they ensure compatibility with distance-based ordination methods and enable faithful geometric representation of ecological relationships. Therefore, understanding and verifying these properties is critical for selecting appropriate measures in downstream beta diversity analyses.

#### 3.2.5 Consequences of Violating Theoretical Conditions

Many commonly used methods—like PCoA, PERMANOVA, and MiRKAT—assume the dissimilarity is Euclidean. But this assumption is often not checked in practice, which can compromise the mathematical validity and reliability of the results. To illustrate the impact of such violations, we present three examples where unmet conditions lead to untrustworthy inferences. Additionally, we introduce diagnostic metrics to quantify the severity of these violations.

##### PCoA

The centered Gram matrix used in PCoA is given by

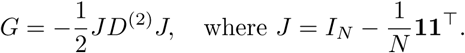

If *D* is Euclidean, then *D*^(2)^ is CND, and the Gram matrix *G* is positive semi-definite (PSD). In this case, all eigenvalues of *G*, denoted by *λ*_1_ ≥ *λ*_2_ ≥ · · · ≥ *λ_N_* ≥ 0, represent non-negative contributions to total variation, and their relative magnitudes describe the proportion of variance captured by each principal coordinate:

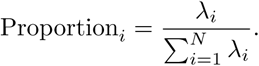

However, when the dissimilarity is non-Euclidean, *G* may contain negative eigenvalues, indicating curvature or non-Euclidean structure in the original space. To quantify this, we define the fraction of negative inertia (FNI) as:

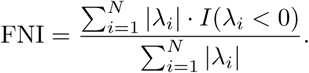

FNI measures the proportion of variation that lies outside of Euclidean space and cannot be faithfully captured by PCoA. In such cases, the standard interpretation of eigenvalues as variance contributions becomes invalid, since the resulting axes no longer represent orthogonal directions in flat Euclidean geometry.

##### PERMANOVA

PERMANOVA evaluates the association between dissimilarity and sample groupings via a pseudo-*F* statistic:

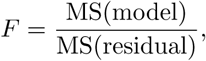

where the mean squares are defined using the Gram matrix *G* and the projection (hat) matrix *H* of the metadata design matrix:

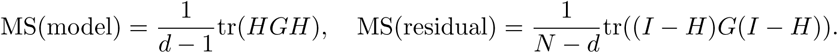

Assuming *G* = *Q*Λ*Q*^⊤^ is the eigen-decomposition of the Gram matrix, we can express:

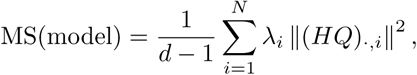

where (*HQ*)_·*,i*_ denotes the projection of the *i*-th principal coordinate onto the metadata space. When *λ_i_ <* 0, the contribution to MS(model) becomes negative, violating the interpretation of variance decomposition and leading to potential misinterpretation of the pseudo-*F* statistic. To address this, we propose a modified pseudo-*F* statistic that considers only positive eigencomponents:

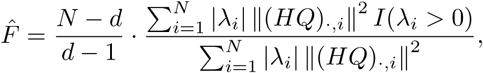

and define the corresponding coefficient of determination as:

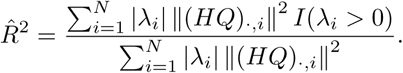

##### MiRKAT

MiRKAT tests the association between a continuous or categorical outcome *Y* and a dissimilarity matrix via a kernel-based score statistic. Let *H* be the projection matrix for confounders and *R_Y_* = (*I* − *H*)*Y* be the residuals. The test statistic is:

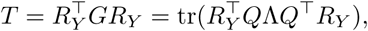

which quantifies the squared association between the outcome residuals and the principal coordinates, weighted by their corresponding eigenvalues. If the dissimilarity is non-Euclidean and some *λ_i_ <* 0, the contribution of those axes becomes negative, undermining the interpretation of *T* as a variance-based measure of association. MiRKAT uses permutation testing to evaluate significance under the null. Given residuals *R*^perm^, the null distribution of the test statistic is approximated by

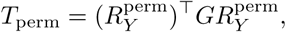

and the *p*-value is computed as:

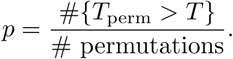

Though this procedure remains valid computationally, the interpretability of *T* as a variance-based statistic is compromised when *G* is not PSD.

##### Summary

These examples demonstrate that when the dissimilarity matrix is not Euclidean, the core assumptions behind commonly used analysis methods can break down. Negative eigenvalues introduce interpretational issues in ordination, hypothesis testing, and variance partitioning. To ensure valid and interpretable results, the use of Euclidean dissimilarities—where the squared dissimilarity matrix is conditionally negative definite—is strongly recommended in downstream beta diversity analysis.

### 3.3 Classification of Dissimilarities

A wide range of dissimilarity measures have been developed to quantify structural differences between microbial communities, each grounded in different mathematical spaces and constructions. However, these measures vary greatly in their theoretical rigor—some are proper metrics, some are Euclidean, and others lack formal geometric properties altogether. While specific dissimilarities such as Bray–Curtis [65] and UniFrac [66, 67] have been studied in isolation, a unified geometric framework is still missing. Here, we summarize the mathematical underpinnings of four common types of dissimilarity measures, based on the metric space in which the community profiles are represented: Euclidean space, Hamming space, the compositional simplex, and the space of probability distributions.

**Euclidean space.** Abundance profiles are often modeled as vectors in R*^K^*, where each coordinate corresponds to a taxon’s count or transformed count. A general class of dissimilarities in this space is based on the Minkowski distance [68]:

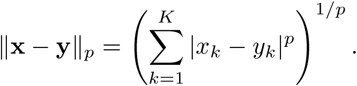

This formulation unifies several widely used distances. For example, *p* = 2 yields the Euclidean distance, while *p* = 1 gives the Manhattan distance. These metrics are compatible with a variety of multivariate techniques—including PCA, PCoA, MANOVA—due to their well-defined geometric properties such as translation invariance, symmetry, and rotational consistency [51, 69–71]. In particular, Euclidean distances preserve variance decomposition and linear structure, making them especially suitable for ordination and hypothesis testing methods discussed in Section 2.2.

**Hamming space.** Presence-absence profiles are naturally embedded in the binary space H*^K^* = {0, 1}*^K^* , where each position indicates whether a taxon is detected [72]. The Hamming distance between two such vectors **p** and **q** is defined as:

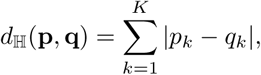

which counts the number of mismatched taxa [72, 73]. Binary similarity measures such as the Jaccard index or unweighted UniFrac are derived from inner products and overlaps in this space.

**Probability space and** *f* **-divergence.** When relative abundances are viewed as discrete probability distributions *P* = (*p*_1_*, . . . , p_K_*), dissimilarity can be quantified using *f* -divergences [74–76], defined as:

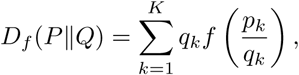

where *f* is a convex function with *f* (1) = 0. Notable examples include the Jensen-Shannon distance [77]

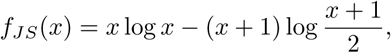

and the Hellinger distance [43, 78]

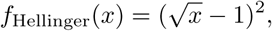

both of which are symmetric and well-suited for measuring shifts in community composition. These divergences provide a statistical interpretation of dissimilarity as information loss or distributional discrepancy.

**Compositional space.** This space is presented last because it is fundamentally different from the other spaces: distances cannot be directly defined within the simplex due to its constrained geometry. Microbial relative abundances are compositional by nature, lying in the simplex:

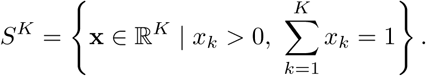

Direct application of Euclidean distance in this constrained space can lead to misleading interpretations [48]. To address this, log-ratio transformations such as the centered log-ratio (CLR) are applied:

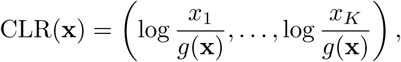

where *g*(**x**) is the geometric mean of the components. The resulting Aitchison distance in the transformed space accounts for the relative scale and compositional nature of microbial communities.

This geometric classification offers a unified view of how dissimilarities map onto different mathematical spaces and provides insights into clustering behavior, ordination suitability, and metric validity. However, it does not explicitly trace the construction of these dissimilarities, from the transformation of single element to the pairwise operations (such as subtraction) performed in geometric spaces. For example, dissimilarities arising from square-root, log, or ratio transformations may share similar analytical behavior despite originating from different geometric spaces.

To address these gaps, we propose a categorization system that classifies dissimilarity measures into four types based on their scaling procedures, data transformations, and the geometric spaces in which distances are computed. This classification helps clarify which dissimilarities are proper metrics, which are Euclidean, and which may pose challenges in downstream analyses such as ordination or hypothesis testing. Table 2 summarizes four major categories. The discussion on the categorizing commonly used dissimilarity measures is detailed in Supplementary Section 2.

**Table 2:**
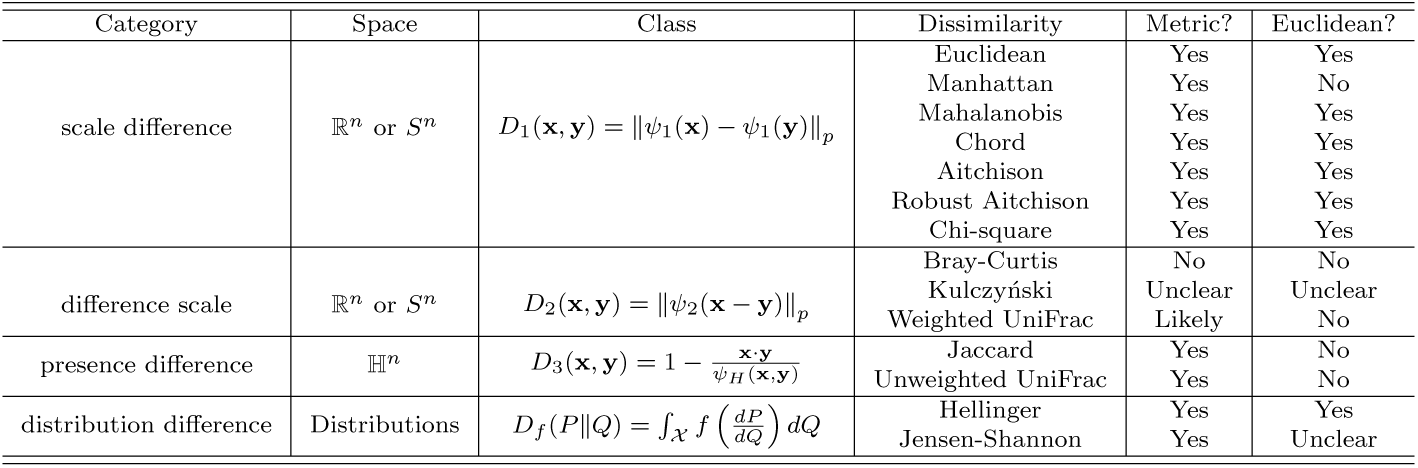
Classification of commonly used dissimilarity measures. Measures of Category 1 operate in Euclidean space (abundance data) or compositional simplex (relative abundance data) and compute the *L_p_* norm after applying a normalization function *ψ*_1_ to each data point individually before calculating the difference. Measures in Category 2 also operate in Euclidean space or compositional simplex but differ by first computing the difference between data points and then applying a normalization function *ψ*_2_. Measures in Category 3 are defined in the Hamming space H*^n^*and usually assess dissimilarity based on presence-absence data. Measures in Category 4 measures the dissimilarity between probability mass functions, which can also be applied to relative abundance data.

**Category 1 (scale difference): Transform-then-Distance (Euclidean embeddings).** Measures in this group apply a normalization function *ψ*_1_(·) to each data point individually (e.g., abundance or relative abundance), and then compute the *L_p_*distance between the transformed vectors:

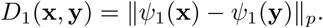

Common examples include Euclidean, Mahalanobis, and Aitchison distances. Notably, when *ψ*_1_(**x**) maps into a Euclidean space (e.g., CLR-transformed compositions), the resulting dissimilarities naturally preserve Euclidean geometry, ensuring compatibility with ordination methods like PCoA and hypothesis tests like PERMANOVA.

**Category 2 (difference scale): Normalize-after-Difference (non-Euclidean forms).** This group computes element-wise differences between communities before normalization:

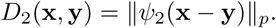

Examples include Bray–Curtis and Weighted UniFrac. These measures often lack a clear geometric embedding and may violate the triangle inequality or the Euclidean property. As a result, distortions can arise in PCoA plots, and negative eigenvalues in the Gram matrix are common. We provide numerical evidence in Section 4.

**Category 3 (presence difference): Presence–absence dissimilarities (Hamming space).** Presence–absence profiles lie in the binary space H*^K^* = {0, 1}*^K^* , where dissimilarity is derived from shared and mismatched entries. Measures like Jaccard and unweighted UniFrac can be written as:

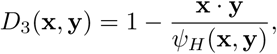

where the denominator counts the number of taxa present in either community. Due to the discrete geometry of Hamming space, these measures typically lack Euclidean embeddability and should be carefully evaluated before downstream analysis.

**Category 4 (distribution difference): Divergence-based dissimilarities (probability space).** Relative abundances can also be viewed as probability mass functions. In this view, *f* -divergences provide a general framework:

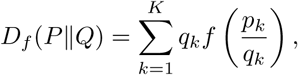

where *f* is a convex function such as the Hellinger or Jensen–Shannon form. These divergences are intuitive and interpretable as average log-ratio deviations, but not all satisfy the triangle inequality or support Euclidean embedding.

### 3.4 Diagnostics and Remedial Corrections

As discussed in Sections 2.1 and 2.2, many widely used dissimilarity measures in microbial ecology lack the Euclidean property, which may lead to negative eigenvalues in the associated Gram matrix and distort downstream analyses such as ordination or hypothesis testing. While structural classifications (Section 2.3) provide useful heuristics, rigorously establishing whether a given dissimilarity is metric or Euclidean remains theoretically nontrivial and often depends on the characteristics of the data [51, 79, 80]. To address this challenge, we develop diagnostic tools for empirically assessing these properties and propose practical remedies for cases where non-Euclidean dissimilarities must be used. The implementations, along with reproducible examples, are available at supplementary material Section 5.

As discussed in Section 2.2, a dissimilarity matrix *D* satisfies the metric property if, for any three distinct samples *i, j, k*, the triangle inequality holds:

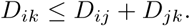

We implement an R function to exhaustively verify this condition across all triplets in the dataset. In addition, we define two sample-level diagnostic scores that quantify the extent to which each sample contributes to collinearity or nonlinearity in the dissimilarity structure:

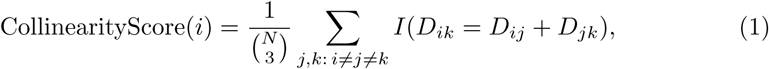

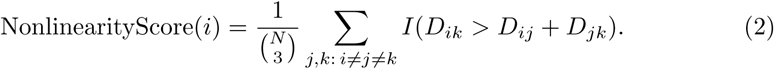

A high collinearity score indicates that the dissimilarity involving sample *i* is exactly additive with respect to other samples, suggesting potential redundancy. In contrast, a high nonlinearity score implies that triangle inequality is violated in configurations involving *i*, suggesting that its information may be poorly captured in methods assuming a Euclidean embedding.

To assess the Euclidean nature of *D*, we construct its centered Gram matrix *G* and perform eigen-decomposition of *G* and compute the FNI introduced in Section 2.2.5. A nonzero FNI indicates that *D* is not Euclidean, and the associated Gram matrix is not positive semi-definite. In such cases, linear dimensionality reduction techniques such as PCoA will introduce geometric distortions, as negative eigenvalues lack interpretable variance decomposition. These diagnostic tools offer a computationally efficient and interpretable framework for assessing the geometric adequacy of dissimilarity matrices, informing the choice of ordination methods and guiding the application of correction procedures when necessary.

When non-Euclidean structure is detected, a natural solution is to select an alternative dissimilarity measure that satisfies Euclidean properties. However, in cases where a non-Euclidean dissimilarity is preferred due to ecological interpretability or established usage, several remedial techniques can be employed to mitigate its limitations in downstream analysis.

One widely used class of remedies involves directly adjusting the dissimilarity matrix to restore metric properties. The ape R package [81] provides two such corrections, namely the Cailliez [82] and Lingoes [83] methods, which aim to remove negative eigenvalues from the centered Gram matrix. The Cailliez method identifies the smallest constant *c* ≥ 0 such that adding *c* uniformly to all off-diagonal entries of the dissimilarity matrix ensures that the resulting Gram matrix becomes positive semi-definite.

This approach follows the analytical solution proposed by Gower and Legendre [51] and implemented in ape. In contrast, the Lingoes method applies a transformation to the squared dissimilarity matrix, adding a constant sufficient to render the matrix conditionally negative definite. Specifically, it transforms the original distances via:

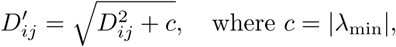

and *λ*_min_ denotes the most negative eigenvalue of the original Gram matrix. This adjustment typically introduces less geometric distortion compared to the Cailliez correction, as it preserves relative relationships among samples more faithfully. Importantly, while both corrections ensure that the modified dissimilarity matrix satisfies the metric condition, they do not guarantee that the resulting distances are strictly Euclidean. That is, negative curvature may still persist in the embedded space, and downstream analyses may continue to be affected by residual non-Euclidean structure. Notably, many widely used microbiome analysis tools—including the ordinate function in the phyloseq package [84]—do not apply these corrections by default. As a result, it remains the user’s responsibility to evaluate and, if needed, preprocess dissimilarity matrices prior to applying ordination methods that assume Euclidean geometry. Similar limitations apply to other analysis pipelines, underscoring the practical relevance of explicit diagnostics and corrections.

When direct modification of the dissimilarity matrix fails to ensure Euclidean structure, an alternative strategy is to adjust the corresponding Gram matrix *G* to enforce positive semidefiniteness (PSD). We introduce two matrix-level remedies: Higham’s nearest PSD approximation [85] and Tikhonov regularization [86].

Higham’s method aims to find the closest PSD matrix *G*^+^ to a given symmetric but indefinite Gram matrix *G*, under the Frobenius norm. The approach begins with the eigen-decomposition *G* = *Q*Λ*Q*^⊤^, where Λ contains the eigenvalues and *Q* the corresponding eigenvectors. Negative eigenvalues in Λ are replaced with zeros to form Λ^+^, and the corrected matrix is reconstructed as *G*^+^ = *Q*Λ^+^*Q*^⊤^. Iterative projections are applied to refine the approximation, ensuring symmetry and numerical stability.

This procedure is widely used in multidimensional scaling and kernel methods, as it guarantees the minimal Frobenius perturbation while restoring the PSD property.

Tikhonov regularization provides an alternative strategy by adding a small ridge term to the diagonal of *G*, yielding the adjusted matrix *G_α_* = *G* + *αI*, where *α >* 0 is a regularization parameter and *I* is the identity matrix. This uniform shift raises all eigenvalues by *α*, rendering *G_α_* PSD when *α* exceeds the absolute value of the most negative eigenvalue. Compared to Higham’s method, Tikhonov regularization minimally distorts the eigenspectrum and preserves the relative contributions of all eigencomponents. However, it modifies the eigenvectors and may introduce rotational distortions in the embedded space.

The choice between these two remedies depends on the analytical goals. Higham’s method strictly enforces non-negativity of eigenvalues and preserves the original eigen-vectors, making it suitable when eigenspace fidelity is essential (e.g., in kernel PCA or spectral clustering). However, by truncating all negative eigenvalues, this approach may lead to a loss of information carried by small but potentially meaningful components [87]. In contrast, Tikhonov regularization maintains the variance distribution across components and avoids abrupt truncation, making it preferable in applications that prioritize smooth eigenvalue decay and numerical conditioning. In practice, we recommend evaluating both methods when correcting non-PSD Gram matrices, and selecting based on their impact on percent variance explained (PVE) and stability in downstream analysis.

Beyond direct corrections to the dissimilarity or Gram matrix, recent methods have been developed to accommodate non-Euclidean structure without enforcing strict Euclidean constraints. Deng et al. [88] proposed Non-Euclidean Multidimensional Scaling (Neuc-MDS), which extends classical MDS by incorporating negative eigenvalues into a generalized bilinear form, allowing sample configurations to be embedded in non-metric spaces while minimizing the STRESS criterion. Gisbrecht and Schleif [89] introduced a scalable strategy for transforming non-metric dissimilarities into positive-definite kernel matrices. Their approach combines Nystroem approximation, double centering, and eigenvalue correction, offering linear computational complexity and suitability for large-scale microbiome datasets.

Together, these strategies provide a flexible toolkit for addressing non-Euclidean dissimilarities in microbial community analysis. Depending on the analytical objective—whether prioritizing ecological interpretability, geometric fidelity, or computational scalability—researchers can select from a range of diagnostic and corrective methods to ensure valid and robust downstream inference.

## 4 Results

### 4.1 Datasets and Analysis Strategy

Building on the theoretical framework in previous sections, we now turn to empirical evaluations across multiple real-world microbiome datasets. Our objective is to examine how often violations of metric and Euclidean properties in commonly used beta diversity measures can be found, and evaluate whether remedial transformations can mitigate such issues. To achieve this objective, we develop a structured evaluation pipeline that quantifies the prevalence of property violations and evaluates their impact on downstream analyses.

1. **Distance Matrix Construction.** For each dataset, we compute pairwise dissimilarities using a diverse panel of beta diversity measures, including presence–absence-based (e.g., Jaccard), abundance-based (e.g., Bray–Curtis), phylogenetically informed (e.g., UniFrac), and compositional (e.g., Aitchison) distances.
2. **Assessment of Metric Violations.** We assess the triangle inequality by evaluating all sample triplets to identify violations and quantify their severity using the number of violations, collinearity, and nonlinearity scores, which provide insight into metric distortions and structural dependencies.
3. **PCoA and Euclidean Properties.** We perform Principal Coordinates Analysis (PCoA) and compute the Fraction of Negative Inertia (FNI), which quantifies the proportion of variance attributed to negative eigenvalues. An FNI of zero indicates that the dissimilarity matrix is fully embeddable in a Euclidean space, while higher values reflect increasing deviation from Euclidean geometry. This provides a diagnostic of how severely a given beta diversity measure violates the Euclidean property and potentially distorts the resulting ordination.
4. **Association via PERMANOVA.** We apply PERMANOVA [90] to assess how well each dissimilarity measure captures compositional variation associated with key covariates. Performance is evaluated using the pseudo-*F* statistic and *R*^2^, which quantify the strength and proportion of group-level separation, respectively. Higher values of pseudo-*F* and *R*^2^ indicate stronger covariate-associated structure, reflecting the informativeness of a dissimilarity measure in detecting biologically meaningful patterns.
5. **Association via MiRKAT.** We apply the MiRKAT framework [19, 25, 91], which uses dissimilarity-induced kernels to test associations with continuous or categorical outcomes. Performance is evaluated using the *R*^2^ statistic, as the permutation-based nature of MiRKAT renders p-values independent of the probability distribution of test statistic and less informative about effect size [92, 93]. Given the typically weak statistical signals in microbiome data, we focus on *R*^2^ as a measure of explanatory power.
6. **Remedial Corrections.** To address non-metric and non-Euclidean distortions, we apply Higham’s matrix correction and Tikhonov regularization. We then re-run PERMANOVA and MiRKAT using the corrected matrices to evaluate improvements in pseudo-*F* and *R*^2^.

We apply the structured analysis pipeline to six diverse microbiome datasets to evaluate the behavior of different beta diversity measures across varied biological contexts.

1. *Cardiometabolic Dataset [94]:* This dataset from the MetaCardis consortium profiles gut microbiome and metabolites in 1,241 middle-aged Europeans across healthy, dysmetabolic, and ischemic heart disease (IHD) groups. Fecal, serum, and urine samples were collected alongside extensive clinical phenotyping to identify cardiometabolic signatures while adjusting for confounders such as polypharmacy and metabolic disorders.
2. *COVID-Gut Microbiome Dataset [95]:* This dataset examines gut microbiome alterations associated with acute SARS-CoV-2 infection and post-recovery states. Conducted at Rutgers University, it includes 60 adults (20 COVID-19 positive, 20 recovered, and 20 healthy controls), with stool samples analyzed by 16S rRNA sequencing. The study identifies sustained depletion of commensal taxa (e.g., Bacteroidaceae, Ruminococcaceae) post-infection, exacerbated by antibiotic use.
3. *IBD Dataset [96, 97]:* This UC-focused dataset integrates metagenomic, proteomic, and metabolomic profiles to investigate microbial protease activity in inflammatory bowel disease. It includes 40 UC patients from UCSD and a validation cohort of 210 individuals (73 UC, 117 Crohn’s disease, 20 healthy controls). The study identifies overabundant Bacteroides vulgatus proteases linked to disease severity.
4. *Pig Gut Microbiome Dataset [98]:* This animal model investigates the impact of long-term fructose and resistant starch intake on gut microbiota and metabolic profiles in Güttingen minipigs. Juvenile pigs were fed high-risk or lower-risk diets over five months, with fecal, plasma, and urine samples analyzed by metagenomics and metabolomics.
5. *ORIGINS Oral Microbiome Dataset [99, 100]:* Part of the ORIGINS study, this dataset evaluates associations between subgingival microbial communities and prediabetes risk. The dataset includes 300 adults aged 20 to 55, with a mean age of 34 years and 77% female participants, from a cohort in the Service Employees International Union 1199. Plaque samples from 1,188 sites were analyzed for 11 periodontitis-related species, alongside clinical periodontal assessments.
6. *HIV-Gut Microbiome Dataset [101]:* This dataset explores the relationship between gut microbiome composition, metabolic disease, and systemic inflammation in people living with HIV (PLWH) and high-risk men who have sex with men (HR-MSM). The dataset includes stool and plasma samples from 113 men, categorized as HIV-negative, untreated HIV-positive, and ART-treated HIV-positive individuals, with further stratification based on lipodystrophy (LD) status. Gut microbiota composition was analyzed using 16S rRNA sequencing, and plasma metabolomics and immune markers were assessed to identify metabolic disease predictors.

### 4.2 Results

Table 3 summarizes the satisfaction of metric properties and FNI of commonly used beta diversity measures across the six datasets (see Supplementary Material Section 4 for additional visualization). Each dissimilarity is evaluated for adherence to the triangle inequality, with “Yes” indicating satisfaction of the metric condition and “No” indicating violations. As expected from theoretical considerations (Table 2), Euclidean, Manhattan, Mahalanobis, Aitchison, and related *L_p_*-based dissimilarities consistently satisfy the metric property across all datasets. Notably, Mahalanobis distance could not be computed for the Cardiometabolic dataset (Dataset (1)) due to singularity in the covariance matrix—a common occurrence when the number of features exceeds the number of samples, as is typical in microbiome data. In contrast, several widely used measures—most notably Bray–Curtis and both versions of weighted UniFrac—frequently violate the triangle inequality, indicating non-metric behavior in specific datasets.

**Table 3:**
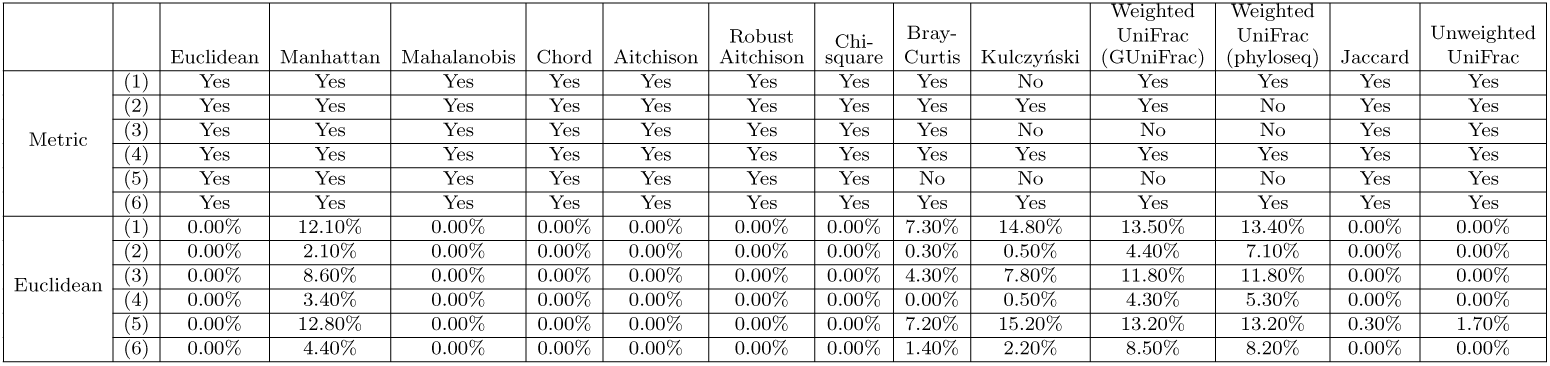
Metric property and Euclidean deviation (FNI) of dissimilarity measures across datasets. The upper block (*Metric*) marks whether each dissimilarity satisfies the metric axioms within each dataset (Yes/No). The lower block (*Euclidean*) reports the Fraction of Negative Inertia (FNI, %) obtained from principal coordinates analysis (PCoA); an FNI of 0% implies the dissimilarity is Euclidean for that dataset, whereas larger values indicate greater deviation from Euclidean geometry. Weighted UniFrac is shown for both GUniFrac and phyloseq implementations. Datasets are indexed (1)–(6).

Bray–Curtis, although widely adopted in microbiome research, fails to satisfy the triangle inequality and is often referred to as a semi-metric [102, 103]. It operates on absolute abundance differences without accounting for compositional constraints or phylogenetic structure, making it susceptible to distortion in high-dimensional microbial data. Our findings confirm its non-metric behavior across all datasets, reinforcing concerns about its use in distance-based analyses. In contrast, Jaccard dissimilarity—a presence-absence–based metric [104–106]—consistently satisfies the triangle inequality. Interestingly, the metric property of weighted UniFrac is software-dependent. The definition implemented in QIIME 2, which follows the formulation in [67], ensures metric properties by omitting the normalization denominator. In contrast, the versions implemented in phyloseq and GUniFrac retain the original form with a denominator [35], leading to systematic violations of the triangle inequality. Our empirical results align with the latter: both phyloseq- and GUniFrac-derived distances consistently fail to meet the metric condition, in contrast to the theoretically valid but less commonly used QIIME 2 variant.

It is important to note that even dissimilarities that qualify as formal metrics (e.g., Jaccard, unweighted UniFrac) may still yield non-Euclidean distance matrices, potentially distorting ordination outcomes. Thus, while metric compliance is a necessary prerequisite for distance-based modeling, it does not guarantee faithful Euclidean embedding—a distinction. Therefore, we also report the Euclidean deviation of each dissimilarity measure using FNI in Table 3. An FNI of zero indicates that the dissimilarity matrix admits an exact Euclidean embedding, whereas higher values signify increasing departures from Euclidean geometry.

As expected, Euclidean distance consistently yields zero FNI across all datasets, as expected from its inner-product–based formulation. This confirms that dissimilarities constructed from *L*_2_ norms preserve Euclidean geometry and can be faithfully represented in PCoA. In contrast, other *L_p_*-based dissimilarities such as Manhattan (*L*_1_) and Mahalanobis—while valid metrics—often yield nonzero FNI values, indicating imperfect Euclidean embeddability. Manhattan distance, in particular, is prone to geometric distortions in high-dimensional microbiome data due to its tendency to produce collinear sample configurations, and Mahalanobis becomes sensitive to ill-conditioned or singular covariance structures. These results underscore that only *L*_2_-based dissimilarities (Chord, Aitchison, Chi-sq) inherently guarantee a Euclidean embedding.

In contrast, Bray–Curtis, which fails the triangle inequality and is formally a semimetric [32, 102], consistently produces nonzero FNI. These distortions are known to affect ordination results [107], introducing artifacts in low-dimensional projections that can mask or exaggerate true ecological gradients [108, 109]. Our results reinforce these concerns by demonstrating that Bray–Curtis not only lacks metric validity but also substantially deviates from Euclidean geometry in empirical data.

Some dissimilarities that satisfy the triangle inequality—such as Jaccard and unweighted UniFrac [15, 67, 104]—also exhibit moderate FNI. This highlights the important distinction between metric and Euclidean properties: being a metric is necessary but not sufficient for guaranteeing a faithful Euclidean representation. For example, Jaccard, while formally metric [105, 106], can be highly sensitive to rare taxa, while unweighted UniFrac is susceptible to exaggerated influence from deep phylogenetic branches [110]. Both can introduce non-Euclidean artifacts in PCoA, despite satisfying metric axioms.

Weighted UniFrac shows the highest and most variable FNI values across datasets. Its implementation is software-dependent: the original form with a normalization denominator [35], used in phyloseq and GUniFrac, fails to satisfy metric properties, while the alternative formulation implemented in QIIME 2 [67] omits the denominator and restores the metric condition. However, even under the latter, the weighted UniFrac remains a tree-based distance and thus inherently non-Euclidean [111, 112]. This explains the persistent presence of negative eigenvalues across all weighted UniFrac variants, emphasizing that correcting for metricity does not guarantee Euclidean behavior, especially for phylogeny-informed distances.

Taken together, these findings underscore the importance of distinguishing between metric validity and Euclidean embeddability when selecting or interpreting dissimilarity measures. While the former ensures theoretical consistency in distance-based modeling, the latter is essential for ordination and kernel-based analyses that rely on inner product geometry.

Figure 2 summarizes the overall structural robustness of each dissimilarity measure using a rose diagram, which aggregates how often a measure satisfies both the metric and Euclidean properties across datasets. A full bar (value of 6) indicates that a dissimilarity consistently behaves as a valid metric and admits a Euclidean embedding, suggesting strong geometric reliability. This integrated view complements the results from Table 3, allowing a more intuitive comparison of robustness across dissimilarity categories.

**Fig. 2:**
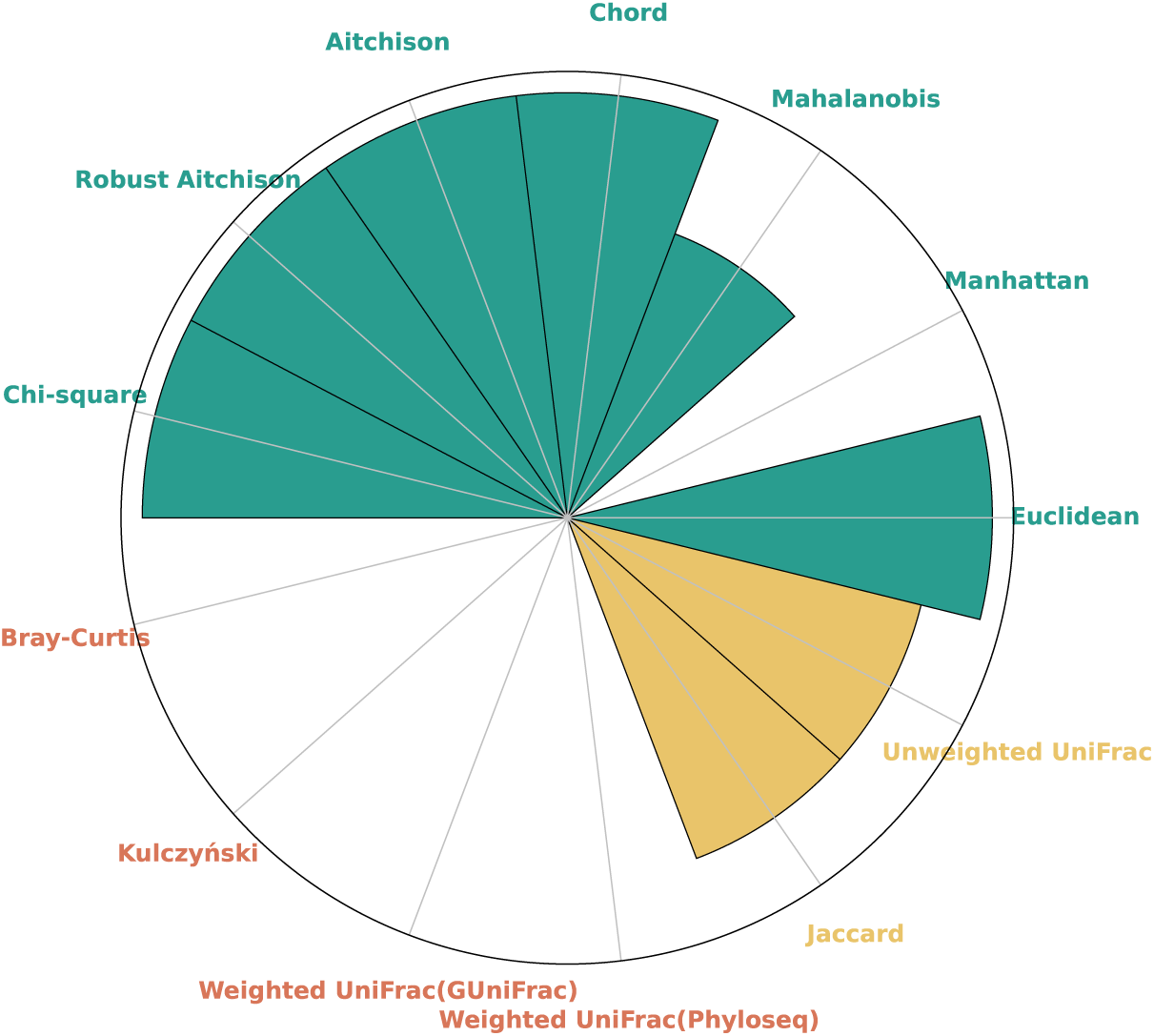
*Frequency of metric and Euclidean behavior across datasets.* This rose diagram summarizes, for each dissimilarity measure, the number of datasets in which it is both metric and Euclidean (i.e., FNI = 0). A full bar (value of 6) indicates consistent metric and Euclidean behavior across all datasets, suggesting greater structural reliability. Bar colors reflect the dissimilarity category.

Overall, dissimilarities based on *L*_2_ norms—such as Euclidean, Chord, Aitchison, Robust Aitchison, and Chi-square—demonstrate the highest structural stability, satisfying both the triangle inequality and Euclidean geometry in nearly all datasets. This is consistent with their theoretical formulations in Table 2, which position them in Category 1. These distances are particularly well suited for ordination and kernel-based inference methods, as they preserve inner-product structure and avoid negative eigenvalues in PCoA. Their robustness also enhances the interpretability of downstream clustering or embedding [107, 108].

In contrast, dissimilarities in Categories 2 and 3 exhibit more variable behavior. Bray–Curtis, although widely used, fails both the metric and Euclidean conditions in all datasets, confirming its semi-metric nature [102, 103]. As noted in prior studies [108, 109], such measures can introduce spurious clustering patterns in compositional datasets, where samples from distinct biological groups may appear artificially close due to projection artifacts. This underscores the risk of using structurally unstable distances in ordination-based microbiome studies. Weighted UniFrac performs particularly poorly in this regard: both the phyloseq and GUniFrac implementations fail to meet metric or Euclidean criteria in any dataset. Even the theoretically valid QIIME 2 variant, while satisfying metric property, remains non-Euclidean due to its dependence on phylogenetic tree structure. This highlights a broader limitation of Category 2 dissimilarities, which may incorporate biologically relevant information but at the cost of geometric interpretability. Interestingly, presence–absence–based measures such as Jaccard and unweighted UniFrac (Category 3) show intermediate robustness. They satisfy the metric condition consistently, but moderate FNI values indicate partial deviation from Euclidean geometry. This reflects a fundamental trade-off: while these measures emphasize rare taxa and phylogenetic breadth, they are more prone to introducing ordination distortions that could obscure true ecological gradients.

Taken together, these findings suggest that the choice of dissimilarity measure should balance biological relevance with geometric stability. While metrics like Bray–Curtis or UniFrac may capture meaningful ecological or phylogenetic signals, their geometric fragility can impair inference in distance-based methods such as PERMANOVA [8] and MiRKAT [19]. Robust dissimilarities—those that maintain both metric and Euclidean properties across datasets—offer greater consistency, reduce the risk of spurious associations, and improve the interpretability of multivariate analyses in microbiome research.

### Comparison of Pseudo F Between No Remedial, Higham Remedial, and Tikhonov Remedial

For dissimilarity matrices that fail to satisfy Euclidean geometry, we apply two remedial correction methods—Higham projection and Tikhonov regularization—and compare their effects on PERMANOVA pseudo-*F* statistics, as shown in Figure 3. The horizontal axis shows pseudo-*F* values computed using uncorrected, non-Euclidean dissimilarities (“No Remedial”), while the vertical axis shows the corresponding values after correction.

**Fig. 3:**
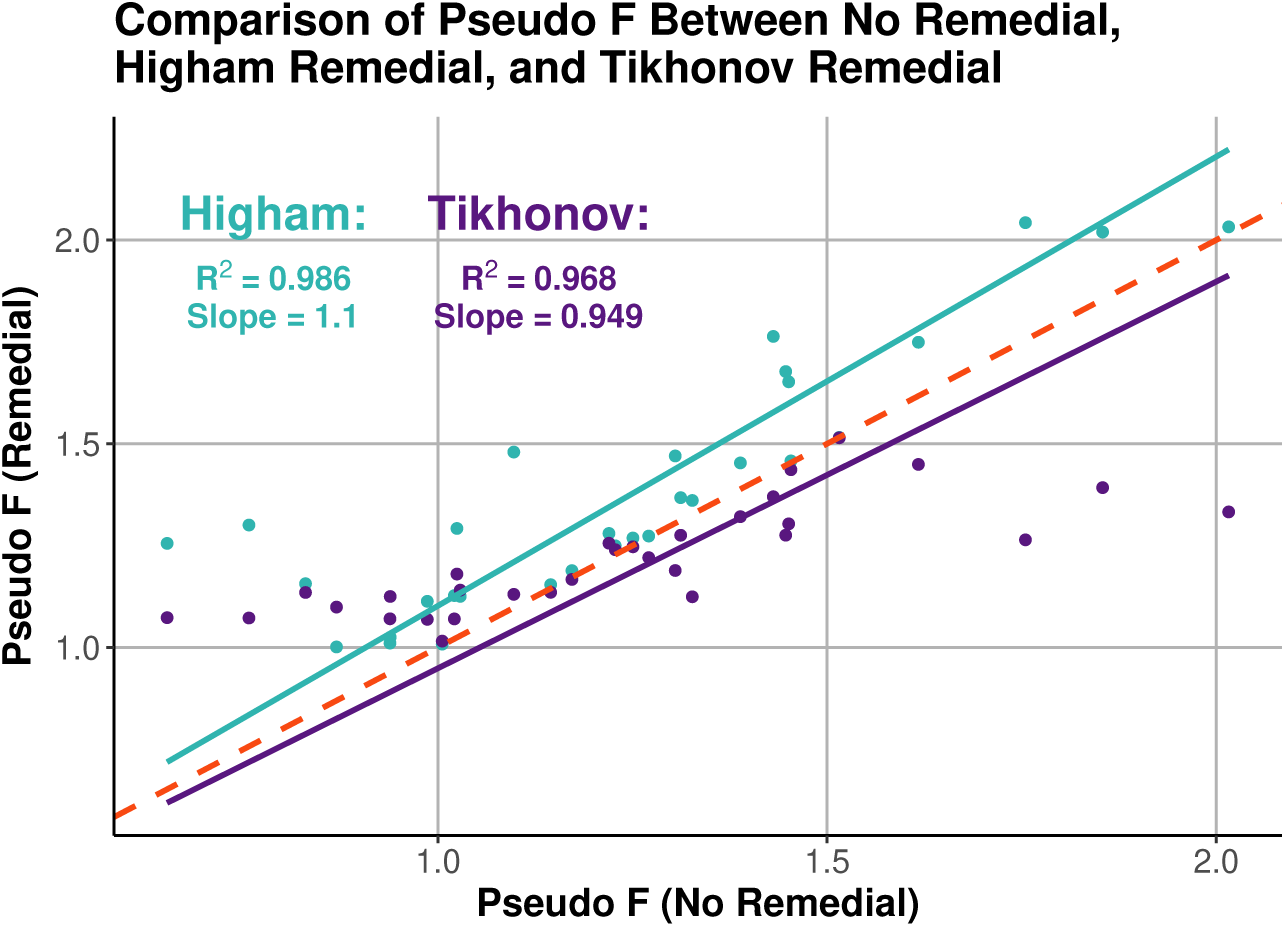
Comparison of PERMANOVA pseudo-F values before and after remedial methods. The horizontal axis shows pseudo-F values obtained using a non-Euclidean dissimilarity measure (“No Remedial”), while the vertical axis shows the corrected pseudo-F values after applying the Higham (teal) or Tikhonov (purple) remedial methods. A red dashed line indicates the 1:1 reference. Regression lines and statistics are shown for each remedial approach; notably, the Higham method increases the original pseudo-F values by roughly 11%.

The Higham method strictly enforces positive semidefiniteness (PSD) by removing all negative eigenvalues from the associated Gram matrix. This results in a systematic inflation of pseudo-*F* values, with a regression slope of 1.10 and *R*^2^ = 0.986, indicating an average 10% increase relative to the original statistics. This consistent upward shift suggests that Higham correction enhances apparent group separability, though it may also exaggerate signal strength due to aggressive spectral truncation. In contrast, the Tikhonov method shifts all eigenvalues uniformly by adding a ridge constant, preserving their relative structure while correcting non-Euclidean behavior. This yields more moderate adjustments to the pseudo-*F* values (slope = 0.949, *R*^2^ = 0.968), reflecting improved numerical stability with less risk of over-amplification.

These results highlight a tradeoff between sensitivity and spectral fidelity: Higham correction may increase statistical power but at the cost of altering the geometry more substantially, whereas Tikhonov offers a more conservative correction. The choice of remedy should therefore be guided by the intended downstream application—e.g., signal detection versus geometry preservation.

Complete results for all assessments—including triangle inequality violations, Euclidean deviation with FNI, and the effects of Higham and Tikhonov remedial corrections on PERMANOVA and MiRKAT—are provided in Supplementary Section 3.

## 5 Discussions and Future Research

In this study, we established a unified mathematical framework for characterizing beta diversity dissimilarities. By systematically analyzing their mathematical formulations, we categorized widely used dissimilarity measures into four structural classes. We further identified three fundamental mathematical properties that an ideal dissimilarity measure should satisfy and analyzed the relationships between these structural categories and the key theoretical properties. This classification framework facilitates intuitive understanding for researchers and practitioners to select appropriate measures in ecological and microbiome studies , and serves as a foundation for the development of new dissimilarity measures with specific desired properties.

To bridge theory and practice, we developed an R-based diagnostic toolkit to assess whether a given dissimilarity matrix satisfies key geometric properties. Our empirical results, drawn from diverse real-world datasets, demonstrate that Euclidean property—more than metric property—is a critical determinant of downstream statistical validity. Notably, the weighted UniFrac distance, though a biologically appealing measure incorporating both abundance and phylogeny, often violates the Euclidean condition and introduces negative eigenvalues in PCoA, undermining its interpretability in multivariate analyses.

Despite these advances, several limitations warrant discussion. First, the scope of datasets used for evaluation remains limited, and the observed violations of Euclidean properties may not generalize across all sample types or environmental contexts. Second, while we provide geometric diagnostics and numerical corrections, the ecological consequences of modifying distance matrices (e.g., via Tikhonov or Higham adjustments) remain to be fully understood. Third, our analysis focused on beta diversity in isolation, without incorporating domain-specific biological factors such as trait structure or spatial autocorrelation.

Future work should explore the robustness of dissimilarity measures to biological confounders such as dominant taxa or rare species inflation. Another promising direction lies in examining how zero-inflated abundance distributions affect dissimilarity-based inference. Since microbiome data often exhibit sparsity and heavy-tailed behavior, it is essential to understand how various measures perform under these realistic constraints. For example, [42] introduced the “double-zero problem,” emphasizing that shared absences between two communities should not contribute to their similarity. Later, [57] argued that dissimilarity values should remain unchanged when common zeros are added to both communities. While these studies highlight the potential impact of zero inflation on dissimilarity measures, a formal mathematical framework to rigorously characterize this effect, quantify the sensitivity of different measures to zero inflation, and assess the effectiveness of proposed remedies is still lacking. Our work provides such a framework and establishes a solid foundation for further investigation.

Furthermore, the integration of beta diversity concepts into multi-omics analysis represents a compelling extension. Recent advances in cross-omics variation modeling, especially in transcriptomics, proteomics, and metabolomics, call for interpretable and geometrically valid dissimilarities to compare sample profiles across omic layers. Beta diversity measures can serve as dissimilarity-based kernels or embedding layers for integrative models, aiding in sample classification, functional alignment, and pathway concordance detection. However, to achieve this goal, it is essential to design dissimilarity measures that not only preserve biological interpretability but also maintain desirable geometric properties across heterogeneous omics domains. Our work lays the foundation for such developments.

## 6 Conclusion

Beta diversity and microbiome data analysis exemplify the kind of interdisciplinary integration that unites biologists, ecologists, mathematicians, and statisticians. Some dissimilarity measures were originally developed to address specific biological questions but lack the mathematical rigor needed for broader analytical reliability. Conversely, mathematically sound distance measures have been adopted to quantify beta diversity, yet they often fall short in capturing biological meaning or meeting practical needs in the field. The sheer number of available metrics has led to widespread—sometimes inappropriate—use, with key assumptions often left unexamined. Paradoxically, the familiarity of certain measures may lead researchers to overlook their theoretical limitations and applicability.

Therefore, we present a geometry-aware framework for the evaluation and application of beta diversity dissimilarity measures. By systematically classifying dissimilarities based on their mathematical formulations and examining their theoretical properties, we identify key structural limitations that compromise the validity of downstream ordination and hypothesis testing. Through diagnostic tools and corrective procedures, we offer practical solutions to address non-Euclidean artifacts and improve inference reliability.

Our findings highlight the importance of aligning ecological interpretation with statistical rigor. In particular, we show that Euclidean compatibility is essential for dissimilarity-based multivariate methods to yield meaningful and robust results. This work not only provides actionable guidance for current microbiome studies but also opens new avenues for integrating beta diversity analysis into cross-omics research and systems biology.

## Data availability

All data supporting the findings of this study are included within the Article and its Supplementary Information files. Raw data of COVID-Gut Microbiome Dataset are publicly available via QIITA (accession 14812). Raw data of HIV-Gut Microbiome Dataset is accessible via the European Bioinformatics Institute (EBI) under accession number ERP125300 and the Metabolomics Workbench under study ID ST001750, subject to institutional agreements and ethical guidelines. The Betadiag software implementing the proposed method—together with the complete scripts and outputs for all real-data analyses—is openly available at supplementary material Section 5.

## Abbreviations

PCoA: Principal Coordinates Analysis
IOS: Illusion of Similarity
CND: Conditional Negative Definiteness
mMDS: Metric Multidimensional Scaling
DPCoA: Double Principal Coordinates Analysis
PSD: Positive Semi-definiteness
FNI: Fraction of Negative Inertia
CLR: Centered Log Ratio
PVE: Percent Variance Explained
Neuc-MDS: Non-Euclidean Multidimensional Scaling

## Declarations

### Ethics approval and consent to participate

Not applicable.

### Consent for publication

Not applicable.

## Authors’ contributions

Z.Z. and Y.Z. conducted the literature review, performed the primary data analysis, developed the computational framework and wrote the initial draft of the manuscript. Z.Z., Y.Z. and W.L. contributed to data preprocessing, simulations, and statistical validation. M.G. provided theoretical guidance on diversity indices, metric properties, and ecological interpretation. L.Z. conceived and supervised the project, guided the statistical methodology and study design, coordinated collaborations across institutions, secured funding and resources, interpreted results in the biomedical context, and rewrote and finalized the manuscript. Z.Z., Y.Z., W.L., M.G., S.S., Y.S., and L.Z. all contributed to manuscript revisions and approved the final version.

## Acknowledgements

We thank Mr. Ruitao Liu from Dr. Liangliang Zhang’s laboratory, Department of Population and Quantitative Health Sciences, School of Medicine, Case Western Reserve University, for preprocessing the raw sequencing data.

## Footnotes

1 A binary function is symmetric if *f* (*x, y*) = *f* (*y, x*).

## References

[1] Costello, E.K., Stagaman, K., Dethlefsen, L., Bohannan, B.J., Relman, D.A.: The application of ecological theory toward an understanding of the human microbiome. Science 336(6086), 1255–1262 (2012)

[2] Gilbert, J.A., Lynch, S.V.: Community ecology as a framework for human microbiome research. Nature medicine 25(6), 884–889 (2019)

[3] Zhu, Y.-G., Zhu, D., Rillig, M.C., Yang, Y., Chu, H., Chen, Q.-L., Penuelas, J., Cui, H.-L., Gillings, M.: Ecosystem microbiome science. Mlife 2(1), 2–10 (2023)

[4] Whittaker, R.H.: Vegetation of the siskiyou mountains, oregon and california. Ecological monographs 30(3), 279–338 (1960)

[5] Koleff, P., Gaston, K.J., Lennon, J.J.: Measuring beta diversity for presence– absence data. Journal of Animal Ecology 72(3), 367–382 (2003)

[6] Pane, C., Sorrentino, R., Scotti, R., Molisso, M., Di Matteo, A., Celano, G., Zaccardelli, M.: Alpha and beta-diversity of microbial communities associated to plant disease suppressive functions of on-farm green composts. Agriculture 10(4), 113 (2020)

[7] Gail, M.H., Wan, Y., Shi, J.: Power of microbiome beta-diversity analyses based on standard reference samples. American Journal of Epidemiology 190(3), 439– 447 (2021)

[8] Anderson, M.J.: Permutational multivariate analysis of variance (permanova). Wiley statsref: statistics reference online, 1–15 (2014)

[9] Real, L.A., Brown, J.H.: Foundations of Ecology: Classic Papers with Commentaries. University of Chicago Press, Chicago (2012)

[10] Turnbaugh, P.J., Ley, R.E., Hamady, M., Fraser-Liggett, C.M., Knight, R., Gordon, J.I.: The human microbiome project. Nature 449(7164), 804–810 (2007)

[11] Wooley, J.C., Godzik, A., Friedberg, I.: A primer on metagenomics. PLoS computational biology 6(2), 1000667 (2010)

[12] Anderson, M.J., Crist, T.O., Chase, J.M., Vellend, M., Inouye, B.D., Freestone, A.L., Sanders, N.J., Cornell, H.V., Comita, L.S., Davies, K.F., et al.: Navigating the multiple meanings of *β* diversity: a roadmap for the practicing ecologist. Ecology letters 14(1), 19–28 (2011)

[13] Jaccard, P.: Etude comparative de la distribution florale dans une portion des alpes et des jura. Bull Soc Vaudoise Sci Nat 37, 547–579 (1901)

[14] Sorensen, T.: A method of establishing groups of equal amplitude in plant sociology based on similarity of species content and its application to analyses of the vegetation on danish commons. Biologiske skrifter 5, 1–34 (1948)

[15] Lozupone, C., Knight, R.: Unifrac: a new phylogenetic method for comparing microbial communities. Applied and environmental microbiology 71(12), 8228– 8235 (2005)

[16] Plantinga, A.M., Wu, M.C.: Beta diversity and distance-based analysis of microbiome data. In: Statistical Analysis of Microbiome Data, pp. 101–127. Springer, New York (2021)

[17] Pasqualini, J., et al.: Charting human gut microbiome states with statistical physics (2025)

[18] Simon, B.O., Nnaji, N.D., Anumudu, C.K., Aleke, J.C., Ekwueme, C.T., Uhegwu, C.C., Ihenetu, F.C., Obioha, P., Ifedinezi, O.V., Ezechukwu, P.S., et al.: Microbiome-based interventions for food safety and environmental health. Applied Sciences 15(9), 5219 (2025)

[19] Zhao, N., Chen, J., Carroll, I.M., Ringel-Kulka, T., Epstein, M.P., Zhou, H., Zhou, J.J., Ringel, Y., Li, H., Wu, M.C.: Testing in microbiome-profiling studies with mirkat, the microbiome regression-based kernel association test. The American Journal of Human Genetics 96(5), 797–807 (2015)

[20] Aboukalam, F., Alharbi, M., Bhatti, M.I.: Improved approximation scales for unreplicated factorial experiments. Journal of Statistical Theory and Applications 21(4), 200–216 (2022)

[21] Chaturvedi, A.: Randomly censored kumaraswamy distribution. Journal of Statistical Theory and Applications 23(1), 1–25 (2024)

[22] Mikhaylov, A., Bhatti, I.M., Dincer, H., Yuksel, S.: Integrated decision recommendation system using iteration-enhanced collaborative filtering, golden cut bipolar for analyzing the risk-based oil market spillovers. Computational Economics 63(1), 305–338 (2024)

[23] Meyners, M., Hasted, A.: On the choice of appropriate models for cata data–a further reply to bi and kuesten. Food Quality and Preference 106, 104818 (2023)

[24] Zhang, Y., Schluter, J., Zhang, L., Cao, X., Jenq, R.R., Feng, H., Haines, J., Zhang, L.: Review and revamp of compositional data transformation: A new framework combining proportion conversion and contrast transformation. Computational and Structural Biotechnology Journal 23, 4088–4107 (2024)

[25] Wilson, N., Zhao, N., Zhan, X., Koh, H., Fu, W., Chen, J., Li, H., Wu, M.C., Plantinga, A.M.: Mirkat: kernel machine regression-based global association tests for the microbiome. Bioinformatics 37(11), 1595–1597 (2021)

[26] Beghini, F., Pullman, J., Alexander, M., Shridhar, S.V., Prinster, D., Singh, A., Matute Júarez, R., Airoldi, E.M., Brito, I.L., Christakis, N.A.: Gut microbiome strain-sharing within isolated village social networks. Nature 637(8044), 167–175 (2025)

[27] Soares, K.O., Da Rocha, T.F., Hale, V.L., Vasconcelos, P.C., Nascimento, L.J., Silva, N.M.V., Rodrigues, A.E., Oliveira, C.J.B.: Comparing the impact of landscape on the gut microbiome of apis mellifera in atlantic forest and caatinga biomes. Scientific Reports 15(1), 5293 (2025)

[28] Raulo, A., Bürkner, P.-C., Finerty, G.E., Dale, J., Hanski, E., English, H.M., Lamberth, C., Firth, J.A., Coulson, T., Knowles, S.C.: Social and environmental transmission spread different sets of gut microbes in wild mice. Nature Ecology & Evolution 8(5), 972–985 (2024)

[29] Kulczyski, S.: Die Pflanzenassoziationen der Pieninen vol. 3. Imprimerie de l’Universit, Lwów (1928)

[30] Léon-Zayas, R., McCargar, M., Drew, J.A., Biddle, J.F.: Microbiomes of fish, sediment and seagrass suggest connectivity of coral reef microbial populations. PeerJ 8, 10026 (2020)

[31] Singh, S., Rinta-Kanto, J.M., Lens, P.N., Kokko, M., Rintala, J., O’Flaherty, V., Ijaz, U.Z., Collins, G.: Microbial community assembly and dynamics in granular, fixed-biofilm and planktonic microbiomes valorizing long-chain fatty acids at 20 c. Bioresource Technology 343, 126098 (2022)

[32] Bray, J.R., Curtis, J.T.: An ordination of the upland forest communities of southern wisconsin. Ecological monographs 27(4), 326–349 (1957)

[33] Pinos, A., Alonso-Alonso, P., Correa-Carmona, Y., Holzmann, K.L., Yon, F., Brehm, G., Steffan-Dewenter, I., Peters, M.K., Weinhold, A., Keller, A.: Host identity, more than elevation, shapes bee microbiomes along a tropical elevation gradient. Frontiers in Microbiology 16, 1671348 (2025)

[34] Kim, Y.-I., Choi, W., Seo, M., Ka, S., Park, J.: Effect of exercise on the human gut microbiota in individuals with overweight and obesity: a systematic review and meta-analysis of randomized controlled trials. Physical Activity and Nutrition 29(2), 49 (2025)

[35] Lozupone, C.A., Hamady, M., Kelley, S.T., Knight, R.: Quantitative and qualitative *β* diversity measures lead to different insights into factors that structure microbial communities. Applied and environmental microbiology 73(5), 1576–1585 (2007)

[36] Fackelmann, G., Manghi, P., Carlino, N., Heidrich, V., Piccinno, G., Ricci, L., Piperni, E., Arrè, A., Bakker, E., Creedon, A.C., et al.: Gut microbiome signatures of vegan, vegetarian and omnivore diets and associated health outcomes across 21,561 individuals. Nature Microbiology 10(1), 41–52 (2025)

[37] Rühlemann, M., Bang, C., Gogarten, J., Hermes, B., Groussin, M., Waschina, S., Poyet, M., Ulrich, M., Akoua-Koffi, C., Deschner, T., et al.: Functional host-specific adaptation of the intestinal microbiome in hominids. Nature communications 15(1), 326 (2024)

[38] Minkowski, H.: Geometrie der Zahlen vol. 1. BG Teubner, Leipzig (1910)

[39] Yang, D., Xu, W.: Clustering on human microbiome sequencing data: a distance-based unsupervised learning model. Microorganisms 8(10), 1612 (2020)

[40] Chen, B., He, X., Pan, B., Zou, X., You, N.: Comparison of beta diversity measures in clustering the high-dimensional microbial data. PloS one 16(2), 0246893 (2021)

[41] Pearson, K.: X. on the criterion that a given system of deviations from the probable in the case of a correlated system of variables is such that it can be reasonably supposed to have arisen from random sampling. The London, Edinburgh, and Dublin Philosophical Magazine and Journal of Science 50(302), 157–175 (1900)

[42] Legendre, P., Gallagher, E.D.: Ecologically meaningful transformations for ordination of species data. Oecologia 129(2), 271–280 (2001)

[43] Hellinger, E.: Neue begründung der theorie quadratischer formen von unendlichvielen veranderlichen. Journal für die reine und angewandte Mathematik 1909(136), 210–271 (1909)

[44] Garcia Mendez, D.F., Egan, S., Wist, J., Holmes, E., Sanabria, J.: Meta-analysis of the microbial diversity cultured in bioreactors simulating the gut microbiome. Microbial ecology 87(1), 57 (2024)

[45] Armetta, J., Li, S.S., Vaaben, T.H., Vazquez-Uribe, R., Sommer, M.O.: Metagenome-guided culturomics for the targeted enrichment of gut microbes. Nature Communications 16(1), 663 (2025)

[46] Mahalanobis, P.C.: On the generalized distance in statistics. Sankhya: The Indian Journal of Statistics, Series A (2008-) 80, 1–7 (2018)

[47] Jayakrishnan, T.T., Sangwan, N., Barot, S.V., Farha, N., Mariam, A., Xiang, S., Aucejo, F., Conces, M., Nair, K.G., Krishnamurthi, S.S., et al.: Multi-omics machine learning to study host-microbiome interactions in early-onset colorectal cancer. NPJ Precision Oncology 8(1), 146 (2024)

[48] Aitchison, J.: The statistical analysis of compositional data. Journal of the Royal Statistical Society: Series B (Methodological) 44(2), 139–160 (1982)

[49] Revel-Muroz, A., Akulinin, M., Shilova, P., Tyakht, A., Klimenko, N.: Stability of human gut microbiome: Comparison of ecological modelling and observational approaches. Computational and Structural Biotechnology Journal 21, 4456–4468 (2023)

[50] Gloor, G.B., Macklaim, J.M., Pawlowsky-Glahn, V., Egozcue, J.J.: Microbiome datasets are compositional: and this is not optional. Frontiers in microbiology 8, 2224 (2017)

[51] Gower, J.C., Legendre, P.: Metric and euclidean properties of dissimilarity coefficients. Journal of classification 3, 5–48 (1986)

[52] Deza, E., Deza, M.M., Deza, M.M., Deza, E.: Encyclopedia of Distances. Springer, New York (2009)

[53] Borg, I., Groenen, P.J.: Modern Multidimensional Scaling: Theory and Applications. Springer, New York (2007)

[54] Whittaker, R.H.: Evolution and measurement of species diversity. Taxon 21(2-3), 213–251 (1972)

[55] Odum, E.P.: The strategy of ecosystem development: An understanding of ecological succession provides a basis for resolving man’s conflict with nature. science 164(3877), 262–270 (1969)

[56] Liberti, L., Lavor, C.: Six mathematical gems from the history of distance geometry. International Transactions in Operational Research 23(3), 897–920 (2016)

[57] Legendre, P., De Cáceres, M.: Beta diversity as the variance of community data: dissimilarity coefficients and partitioning. Ecology Letters 16(8), 951–963 (2013)

[58] Rao, C.R.: Diversity and dissimilarity coefficients: a unified approach. Theoretical population biology 21(1), 24–43 (1982)

[59] Anderson, M.J.: Distance-based tests for homogeneity of multivariate dispersions. Biometrics 62(1), 245–253 (2006)

[60] Pavoine, S., Dufour, A.-B., Chessel, D.: From dissimilarities among species to dissimilarities among communities: a double principal coordinate analysis. Journal of theoretical biology 228(4), 523–537 (2004)

[61] Menger, K.: Untersuchungen über allgemeine metrik. Mathematische Annalen 100, 75–163 (1928)

[62] Blumenthal, L.M.: A modern view of distance geometry. Bulletin of the American Mathematical Society 50(1), 6–16 (1944)

[63] Blumenthal, L.M.: Theory and Applications of Distance Geometry. Oxford University Press, Oxford, UK (1953)

[64] Gower, J.C.: Properties of euclidean and non-euclidean distance matrices. Linear algebra and its applications 67, 81–97 (1985)

[65] Greenacre, M., Primicerio, R.: Chapter: Measures of distance between samples: Non-euclidean. Universitat Pompeu Fabra Barcelona Department of Economics and Business [Online]. http://www.econ.up.edu/michael/stanford/maeb5.pdf (2008)

[66] Schloss, P.D.: Evaluating different approaches that test whether microbial communities have the same structure. The ISME journal 2(3), 265–275 (2008)

[67] Lozupone, C., Lladser, M.E., Knights, D., Stombaugh, J., Knight, R.: Unifrac: an effective distance metric for microbial community comparison. The ISME journal 5(2), 169–172 (2011)

[68] Conway, J.B.: A Course in Functional Analysis vol. 96. Springer, New York (2019)

[69] Seal, H.: Multivariate statistical analysis for biologists. (1964)

[70] DasGupta, A.: Probability for Statistics and Machine Learning: Fundamentals and Advanced Topics. Springer, New York (2011)

[71] Mardia, K.V., Kent, J.T., Taylor, C.C.: Multivariate Analysis. John Wiley & Sons, Hoboken, NJ, USA (2024)

[72] Ash, R.B.: Information Theory. Courier Corporation, Boston, MA (2012)

[73] Richardson, T., Urbanke, R.: Modern Coding Theory. Cambridge university press, Cambridge, UK (2008)

[74] Haroutunian, M., Mkhitaryan, K., Mothe, J.: f-divergence measures for evaluation in community detection. In: Proceedings of the Workshop on Collaborative Technologies and Data Science in Smart City Applications, pp. 1–6 (2018). 10.1145/3289256.3289260

[75] Nishiyama, T.: Generalized bregman and jensen divergences which include some f-divergences. arXiv preprint arXiv:1808.06148 (2018)

[76] Wainwright, M.J.: High-Dimensional Statistics: A Non-Asymptotic Viewpoint. Cambridge University Press, Cambridge, UK (2019). https://www.cambridge.org/core/books/highdimensional-statistics/8A91ECEEC38F46DAB53E9FF8757C7A4E

[77] Lin, J.: Divergence measures based on the shannon entropy. IEEE Transactions on Information Theory 37(1), 145–151 (1991)

[78] Beran, R.: Minimum hellinger distance estimates for parametric models. The Annals of Statistics 5(3), 445–463 (1977) 10.1214/aos/1176343842

[79] Greenberg, M.J.: Euclidean and non-Euclidean Geometries: Development and History. Macmillan, New York, NY, USA (1993)

[80] De Santis, E., Martino, A., Rizzi, A.: On component-wise dissimilarity measures and metric properties in pattern recognition. PeerJ Computer Science 8, 1106 (2022)

[81] Paradis, E., Schliep, K.: ape 5.0: an environment for modern phylogenetics and evolutionary analyses in r. Bioinformatics 35(3), 526–528 (2019)

[82] Cailliez, F.: The analytical solution of the additive constant problem. Psychometrika 48(2), 305–308 (1983)

[83] Lingoes, J.C.: Some boundary conditions for a monotone analysis of symmetric matrices. Psychometrika 36(2), 195–203 (1971)

[84] McMurdie, P.J., Holmes, S.: phyloseq: an r package for reproducible interactive analysis and graphics of microbiome census data. PloS one 8(4), 61217 (2013)

[85] Higham, N.J.: Computing a nearest symmetric positive semidefinite matrix. Linear Algebra and its Applications 103, 103–118 (1988) 10.1016/0024-3795(88)90223-6

[86] Tikhonov, A.N.: Solution of incorrectly formulated problems and the regularization method. Soviet Mathematics Doklady 4, 1035–1038 (1963)

[87] Legendre, P.: Numerical Ecology. Elsevier, Amsterdam (2012)

[88] Deng, C., Gao, J., Lu, K., Luo, F., Sun, H., Xin, C.: Neuc-mds: Non-euclidean multidimensional scaling through bilinear forms. Advances in Neural Information Processing Systems 37, 121539–121569 (2024)

[89] Gisbrecht, A., Schleif, F.-M.: Metric and non-metric proximity transformations at linear costs. Neurocomputing 167, 643–657 (2015)

[90] Anderson, M.J.: A new method for non-parametric multivariate analysis of variance. Austral ecology 26(1), 32–46 (2001)

[91] Jiang, Z., He, M., Chen, J., Zhao, N., Zhan, X.: Mirkat-mc: a distance-based microbiome kernel association test with multi-categorical outcomes. Frontiers in Genetics 13, 841764 (2022)

[92] Pesarin, F., Salmaso, L.: Permutation Tests for Complex Data: Theory, Applications and Software. John Wiley & Sons, Hoboken, NJ (2010)

[93] Edgington, E., Onghena, P.: Randomization Tests. Chapman and Hall/CRC, Boca Raton, FL, USA (2007)

[94] Fromentin, S., Forslund, S.K., Chechi, K., Aron-Wisnewsky, J., Chakaroun, R., Nielsen, T., Tremaroli, V., Ji, B., Prifti, E., Myridakis, A., et al.: Microbiome and metabolome features of the cardiometabolic disease spectrum. Nature medicine 28(2), 303–314 (2022)

[95] Yin, Y.S., Minacapelli, C.D., Parmar, V., Catalano, C.C., Bhurwal, A., Gupta, K., Rustgi, V.K., Blaser, M.J.: Alterations of the fecal microbiota in relation to acute covid-19 infection and recovery. Molecular biomedicine 3(1), 36 (2022)

[96] Gonzalez, C.G., Mills, R.H., Zhu, Q., Sauceda, C., Knight, R., Dulai, P.S., Gonzalez, D.J.: Location-specific signatures of crohn’s disease at a multi-omics scale. Microbiome 10(1), 133 (2022)

[97] Mills, R.H., Dulai, P.S., Vázquez-Baeza, Y., Sauceda, C., Daniel, N., Gerner, R.R., Batachari, L.E., Malfavon, M., Zhu, Q., Weldon, K., et al.: Multi-omics analyses of the ulcerative colitis gut microbiome link bacteroides vulgatus proteases with disease severity. Nature microbiology 7(2), 262–276 (2022)

[98] Curtasu, M.V., Tafintseva, V., Bendiks, Z.A., Marco, M.L., Kohler, A., Xu, Y., Nørskov, N.P., Nygaard Lærke, H., Bach Knudsen, K.E., Hedemann, M.S.: Obesity-related metabolome and gut microbiota profiles of juvenile güttingen minipigs—long-term intake of fructose and resistant starch. Metabolites 10(11), 456 (2020)

[99] Demmer, R., Jacobs Jr, D., Singh, R., Zuk, A., Rosenbaum, M., Papapanou, P., Desvarieux, M.: Periodontal bacteria and prediabetes prevalence in origins: the oral infections, glucose intolerance, and insulin resistance study. Journal of dental research 94(9 suppl), 201–211 (2015)

[100] Demmer, R.T., Breskin, A., Rosenbaum, M., Zuk, A., LeDuc, C., Leibel, R., Paster, B., Desvarieux, M., Jacobs Jr, D.R., Papapanou, P.N.: The sub-gingival microbiome, systemic inflammation and insulin resistance: the oral infections, glucose intolerance and insulin resistance study. Journal of clinical periodontology 44(3), 255–265 (2017)

[101] Armstrong, A.J., Quinn, K., Li, S.X., Schneider, J.M., Nusbacher, N.M., Doenges, K.A., Fiorillo, S., Marden, T.J., Higgins, J., Reisdorph, N., et al.: Systems analysis of gut microbiome influence on metabolic disease in hiv and high-risk populations. bioRxiv, 2021–03 (2021)

[102] Legendre, P., Legendre, L., et al.: Numerical ecology: developments in environmental modelling. Developments in Environmental Modelling 20(1) (1998)

[103] Gromov, M., Katz, M., Pansu, P., Semmes, S.: Metric Structures for Riemannian and non-Riemannian Spaces vol. 152. Springer, New York (1999)

[104] Jaccard, P.: Nouvelles recherches sur la distribution florale. Bull. Soc. Vaud. Sci. Nat. 44, 223–270 (1908)

[105] Lipkus, A.H.: A proof of the triangle inequality for the tanimoto distance. Journal of Mathematical Chemistry 26(1), 263–265 (1999)

[106] Kosub, S.: A note on the triangle inequality for the jaccard distance. Pattern Recognition Letters 120, 36–38 (2019)

[107] Gower, J.C.: Some distance properties of latent root and vector methods used in multivariate analysis. Biometrika 53(3-4), 325–338 (1966)

[108] Pawlowsky-Glahn, V., Egozcue, J.J.: Compositional data and their analysis: an introduction. Geological Society, London, Special Publications 264(1), 1–10 (2006)

[109] Quinn, T.P., Erb, I., Richardson, M.F., Crowley, T.M.: Understanding sequencing data as compositions: an outlook and review. Bioinformatics 34(16), 2870–2878 (2018)

[110] Barwell, L.J., Isaac, N.J., Kunin, W.E.: Measuring *β*-diversity with species abundance data. Journal of Animal Ecology 84(4), 1112–1122 (2015)

[111] Gupta, A.: Embedding tree metrics into low dimensional euclidean spaces. In: Proceedings of the Thirty-first Annual ACM Symposium on Theory of Computing, pp. 694–700 (1999)

[112] Chen, J., Safro, I.: Algebraic distance on graphs. SIAM Journal on Scientific Computing 33(6), 3468–3490 (2011)

