## Supplementary for "Mathematical Foundations of Beta Diversity: Why Common Metrics Fail in Microbiome Analysis"

#### 1 A Simple Bibliometric Study

The modern framework distinguishing alpha, beta, and gamma diversity was formally introduced by Robert H. Whittaker in the 1960s [1], with the goal of describing how biodiversity is partitioned across environmental or spatial gradients. In this framework, alpha diversity refers to the species diversity within a single site or local habitat, while gamma diversity refers to the total species diversity across different habitats within a large geographic area. The connection between these scales is mediated by beta diversity, which characterizes the turnover or differentiation between local communities [2, 3]. Whittaker's work laid the foundation for using dissimilarity measures in beta diversity analysis, allowing ecologists to better understand the complexity of ecological communities.

In the subsequent decades, the concept of beta diversity has expanded beyond its origins in plant ecology to become a cornerstone of biodiversity research in diverse ecosystems [4, 5]. A growing recognition that community composition can vary significantly even within seemingly homogeneous environments has driven interest in more granular analyses of beta diversity. At the same time, advances in statistical and computational methodologies, alongside the rise of high-throughput sequencing technologies, have enabled researchers to explore ecological variation at unprecedented scales. These developments have facilitated macroecological studies of forests and coral reefs [6] as well as fine-scale investigations of microbial communities in soils, oceans, and host-associated environments [7, 8]. In microbiome research, where communities are inherently dynamic and often shaped by host physiology, environmental gradients, and evolutionary history, beta diversity plays a crucial role in elucidating patterns of microbial variation and their functional implications.

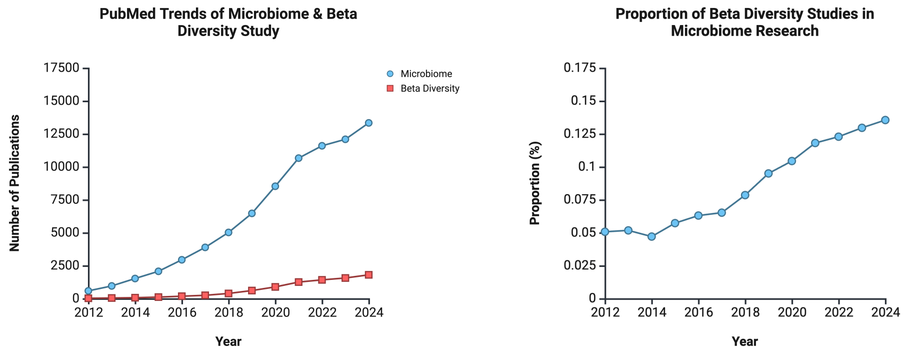


**Fig. 1** Upper: The number of publications mentioning "microbiome" (blue) and both "microbiome" and "beta diversity" (red) indexed in PubMed from 2012 to 2024. Microbiome-related studies have grown exponentially, exceeding 10,000 publications per year, while beta diversity studies have shown a steady increase. Lower: The proportion of beta diversity studies within microbiome research, which has more than doubled from approximately 5% in 2012 to over 12% in 2024, reflecting the increasing recognition of beta diversity in microbial ecology. Analysis and visualization are done using R package `RISmed` and `ggplot2`.

The increasing prominence of beta diversity in microbiome research is reflected in publication trends. A bibliometric analysis of PubMed-indexed studies (Figure 1) shows that publications on microbiome research have experienced exponential growth over the past decade, surpassing 10,000 papers per year in recent years. While beta diversity studies constitute a smaller fraction of this body of literature, they have also shown a steady upward trajectory, with their proportion among microbiome-related studies more than doubling from 5% to over 12% since 2012 (Figure 1). This trend underscores the growing recognition of beta diversity as a fundamental concept in microbial ecology and suggests that understanding compositional variation is becoming increasingly central to microbiome research. Despite this expansion, however, there remains a lack of consensus on how best to quantify beta diversity in different microbiome contexts.

### 2 Introduction of Commonly Used Beta Diversity Measures

Let $N$ denote the number of samples and $K$ the number of observed taxa. For sample $i$, let $a_{ik}$ represent the absolute abundance (sequence read counts) of taxon $k$, with corresponding relative abundance $r_{ik} = a_{ik}/a_{i\cdot}$, where $a_{i\cdot} = \sum_{k=1}^{K} a_{ik}$. Let $p_{ik} = I(a_{ik} > 0)$ denote presence–absence indicators.

#### 2.1 Jaccard Distance

The Jaccard index, introduced by the Swiss botanist Paul Jaccard in 1901 [9], is one of the earliest measures of similarity and diversity between two sets of species. Originally, it considered the presence-absence information of the elements in two finite sets $A$ and $B$. The Jaccard similarity index is mathematically defined as:

$$J(A, B) = \frac{|A \cap B|}{|A \cup B|}.$$

Here $|\cdot|$ is the cardinality of a set. Hence, $|A \cap B|$ represents the number of elements present in both $A$ and $B$ and $|A \cup B|$ represents the number of elements present in $A$ or $B$. The Jaccard index yields a value between 0 and 1, where 0 denotes disjoint sets and 1 denotes identical sets. The corresponding Jaccard distance, often used as a dissimilarity measure, is given by:

$$d_J(A, B) = 1 - J(A, B) = 1 - \frac{|A \cap B|}{|A \cup B|}.$$

This definition yields a proper distance-like metric that increases as the sets become less similar and satisfies the properties of a metric on the space of all finite sets [10–12]. For presence–absence data, the Jaccard distance between samples $i$ and $j$ is defined as

$$d_J(i, j) = 1 - \frac{\sum_k p_{ik} p_{jk}}{\sum_k I(p_{ik} + p_{jk} > 0)}.$$

The Jaccard index extends to non-binary data by representing sets as non-negative vectors, denoting degrees of membership. In this setting, set operations are replaced with element-wise minima and maxima. For abundance data, the generalized (non-binary) Jaccard similarity index [13] is defined as

$$J_{\text{non-binary}}(i,j) = \frac{\sum_k \min(a_{ik}, a_{jk})}{\sum_k \max(a_{ik}, a_{jk})}.$$

This extension accommodates continuous or abundance data, allowing for a more nuanced comparison of communities or feature sets. The non-binary Jaccard index collapses to the binary case when all values are restricted to 0 or 1. After more than a century, Jaccard-based measures remain indispensable in a broad range of applications, capturing essential notions of similarity and difference with a conceptual elegance that has proven timeless. It continues to find widespread application in fields such as ecology, information retrieval, and machine learning [14].

#### 2.2 Kulczyński Dissimilarity

The Kulczyński dissimilarity is named after Stanisław Kulczyński, a Polish botanist and phytosociologist. In 1928, Kulczyński introduced this measure to quantify the similarity between ecological communities, originally using species presence-absence data [15]. His work focused on understanding the relationships among vegetation samples, and the Kulczyński dissimilarity has since become a valuable tool in studies of ecology, community structure, and biodiversity assessment.

The original binary version based on presence-absence data is given by:

$$S_{ij}^{(\text{binary})} = \frac{1}{2}\left(\frac{\sum_k p_{ik}p_{jk}}{\sum_k p_{ik}} + \frac{\sum_k p_{ik}p_{jk}}{\sum_k p_{jk}}\right).$$

This formula reflects the original form of the measure, which accounts purely for species occurrences without considering their abundances. The definition was later extended to handle species abundance, making it a more nuanced measure of dissimilarity. This extension is especially useful when in contexts where the proportional overlap of species abundances between samples is of interest.

For abundance data, the Kulczyński similarity index between two samples $i$ and $j$ is defined as [16–18]:

$$S_{ij} = \frac{1}{2}\left(\frac{\sum_k \min(a_{ik},\, a_{jk})}{a_{i\cdot}} + \frac{\sum_k \min(a_{ik},\, a_{jk})}{a_{j\cdot}}\right).$$

The term $\sum_k \min(a_{ik},\, a_{jk})$ captures the shared abundance of each species between the two samples, highlighting overlap in species composition. The denominators $a_{i\cdot} = \sum_k a_{ik}$ and $a_{j\cdot} = \sum_k a_{jk}$ normalize the shared abundances relative to the overall abundances. The Kulczyński dissimilarity is then calculated by subtracting the similarity from one [19]:

$$D_{ij} = 1 - S_{ij} = \frac{1}{2}\left(\frac{\sum_k |a_{ik} - a_{jk}|}{a_{i\cdot}} + \frac{\sum_k |a_{ik} - a_{jk}|}{a_{j\cdot}}\right).$$

By averaging the normalized shared abundances and subtracting from one, the Kulczyński dissimilarity balances contributions from both samples and provides a measure of dissimilarity that does not overly penalize species with small distribution ranges. It is also considered a suitable alternative to the Jaccard distance for analyzing groups of species with overlapping distributions, as species with small distribution areas can be grouped alongside those with larger ones when there is a subset relationship [20–22]. This makes the Kulczyński distance a valuable tool in ecological studies aiming to assess biodiversity patterns and community structures across different environments or temporal scales.

#### 2.3 Bray-Curtis Dissimilarity

The Bray–Curtis dissimilarity was first introduced by Bray and Curtis [23] in 1957. They developed this measure to interpret complex ecological data related to forest communities. Since then, the Bray–Curtis dissimilarity has become one of the most well-known tools to quantify compositional differences among ecosystems. It assesses the difference between two communities via the abundances of various species present in each community. Its modern form is mathematically defined as

$$\mathrm{BC}_{ij} = \frac{\sum_k |a_{ik} - a_{jk}|}{\sum_k (a_{ik} + a_{jk})}.$$

This formulation quantifies the dissimilarity by summing the absolute differences in the abundances of species and normalizing it by the total abundances of both communities. In practice, if relative abundances are used for this calculation, Bray-Curtis dissimilarity is just half the Manhattan distance as shown in Supplementary Equation (1). Notably, the Bray–Curtis dissimilarity is also mathematically related to the quantitative Sørensen similarity index [24, 25] $\mathrm{QS}_{ij}$, as both emphasize shared species abundance while ignoring joint absences. The formula can be expressed as

$$\mathrm{BC}_{ij} = 1 - \mathrm{QS}_{ij} = 1 - \frac{2\sum_k \min(a_{ik},\ a_{jk})}{\sum_k (a_{ik} + a_{jk})}.$$

However, it is important to note that the Bray–Curtis dissimilarity cannot be considered as a true distance because it does not satisfy the triangle inequality [26]. Furthermore, as [27] pointed out, Bray-Curtis dissimilarity between two communities often comes from the difference in "size" (or sequencing depth) rather than true composition. These limitations mean that while the Bray–Curtis dissimilarity is highly effective for comparing community compositions, it may not be suitable for analysis that rely on the properties of distance, such as certain ordination techniques and distance-based clustering algorithms.

#### 2.4 UniFrac Dissimilarity

Unlike traditional ecological diversity metrics designed for plants and animals, UniFrac dissimilarity was specifically developed for microbiome research [28]. By incorporating phylogenetic relationships among microbial taxa, it refines community similarity assessment beyond what abundance- or presence-based measures offer. Both unweighted and weighted versions of UniFrac will be introduced in the following paragraphs.

The unweighted UniFrac distance was developed by Lozupone and Knight [29] in 2005. It quantifies the dissimilarity between microbial communities by leveraging phylogenetic information, focusing exclusively on the prevalence of taxa [29]. Mathematically, it is defined as the fraction of the lengths of unshared branches for two communities in a phylogenetic tree, that is,

$$d_{\text{Unweighted UniFrac}}(i,j) = \frac{\sum_{k=1}^{\bar{K}} b_k |p_{ik} - p_{jk}|}{\sum_{k=1}^{\bar{K}} b_k I(p_{ik} + p_{jk} > 0)},$$

where $\bar{K}$ denotes the number of branches and $b_k$ is the branch length of the $k$-th edge. The unweighted UniFrac dissimilarity accounts for evolutionary relationships among taxa, offering a robust approach to capture community differences beyond simple taxonomic comparisons. Considering the presence or absence of taxa, it emphasizes rare or unique evolutionary lineages, which makes it particularly valuable in ecological studies where such lineages might reflect critical environmental adaptations or evolutionary pressures [29–31]. A graph illustration on calculating the unweighted UniFrac distance between two samples is provided in Figure 2.

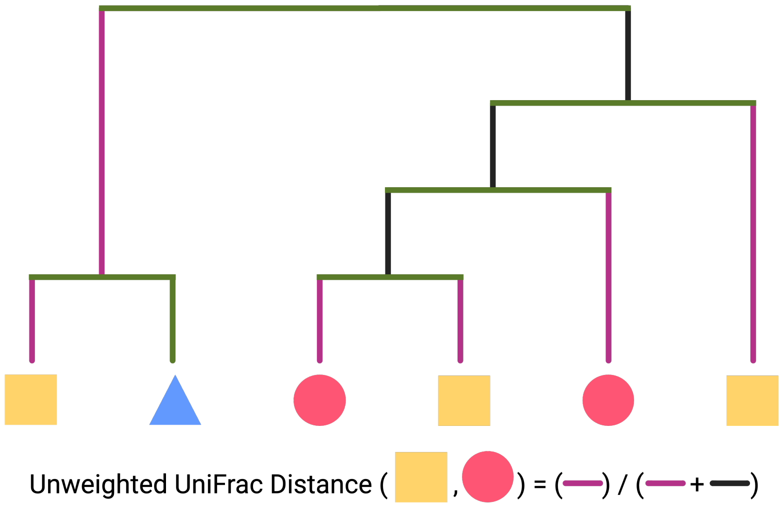


**Fig. 2 Unweighted UniFrac between two samples 'Circle' and 'Square'.** The blue triangle represents a taxon that does not belong to either of the two samples. The unshared branch lengths, representing phylogenetic branches unique to each community (highlighted in purple), are summed and divided by the sum of the unshared branch lengths and shared branch lengths (highlighted in black). This distance quantifies the phylogenetic dissimilarity between the two communities by comparing unique evolutionary lineages to the overall tree structure.

The Weighted UniFrac, introduced by Lozupone et al. [32] in 2007, extends the unweighted UniFrac distance by incorporating the relative abundances of taxa within microbial communities. Unlike the unweighted version, weighted UniFrac quantifies the dissimilarity between communities not only based on the phylogenetic distances of taxa but also by weighting these distances according to their relative abundances [32, 33]. Mathematically, it is defined as [32]:

$$d_{\text{Weighted UniFrac}}(i,j) = \frac{\sum_{k=1}^{\bar{K}} b_k |a_{ik} - a_{jk}|}{\sum_{k=1}^{\bar{K}} b_k |a_{ik} + a_{jk}|}$$

The numerator accounts for the weighted differences in abundance along each branch, while the denominator normalizes this difference by the total branch length of the phylogenetic tree. By combining evolutionary relationships and relative abundances, Weighted UniFrac offers a more nuanced measure of community dissimilarity, capturing both taxonomic composition and ecological dominance. As a result, it particularly useful in studies where abundance disparities between communities are important, such as those involving environmental gradients or host-associated [28, 32].

#### 2.5 Minkowski Distance

Minkowski distances, introduced by Hermann Minkowski in the early 20th century [34], form a family of metrics that unify various notions of distance under a single mathematical framework. For points $x = (x_1, x_2, \ldots, x_K)$ and $y = (y_1, y_2, \ldots, y_K)$ in a $K$-dimensional space, the Minkowski distance is defined as

$$d_p(x,y) = \left( \sum_{i=1}^{K} |x_i - y_i|^p \right)^{1/p},$$

where $p \geq 1$. For microbiome data, the Minkowski distance between community $i$ and $j$ is

$$d_p(i,j) = \left( \sum_{k=1}^{K} |a_{ik} - a_{jk}|^p \right)^{1/p}.$$

The Minkowski distances qualify as true distance ensured by the Minkowski inequality [35]. By adjusting the parameter $p$, one can seamlessly transition between different distance measures, each characterized by distinct geometric and analytical properties.

When $p = 2$, the Minkowski distance becomes a special case known as the Euclidean distance

$$d_2(x,y) = \sqrt{\sum_{i=1}^{K} (x_i - y_i)^2},$$

whose roots trace back to Euclid's Elements [36]. The Euclidean distance represents the shortest straight-line path between two points, making it intuitively appealing and widely used. A key property of Euclidean distance is its rotational invariance: rotations and translations of the coordinate system do not alter the measured distances. This

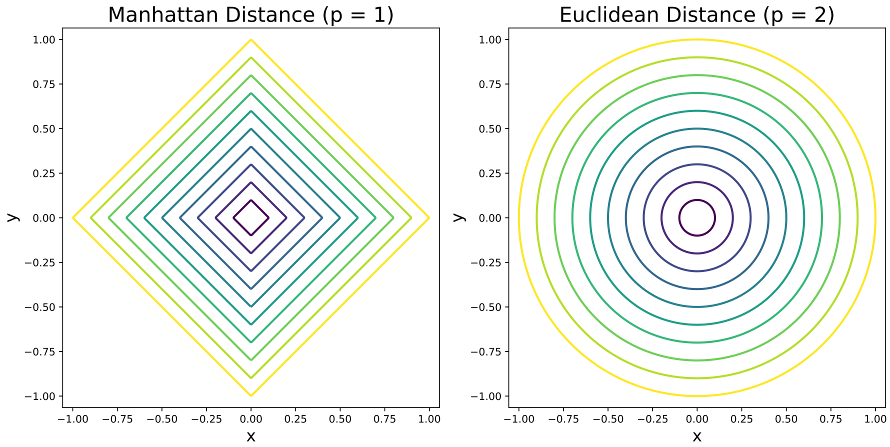


**Fig. 3** Comparison of level sets for Manhattan distance ($p = 1$) and Euclidean distance ($p = 2$), illustrating the diamond-shaped geometry of Manhattan distance and the circular geometry of Euclidean distance.

invariance is crucial in many applications, ensuring that the distance captures intrinsic relationships rather than coordinate-based artifacts. As a result, Euclidean distance serves as the foundation for numerous statistical and machine learning techniques, including principal component analysis [37–39], clustering methods like K-means [40, 41], and regularized linear regression [42, 43]. Its squared form also appears naturally in optimization problems, where minimizing squared Euclidean distances often simplifies to solvable analytical solutions [44].

When $p = 1$, the Minkowski distance becomes a special case known as the Manhattan (or taxicab) distance

$$d_1(x, y) = \sum_{i=1}^{K} |x_i - y_i|.$$

It is so named because it reflects distances one might travel along a grid of perpendicular city streets [45]. Unlike the Euclidean distance, the Manhattan distance does not directly measure "straight-line" displacement; instead, it sums absolute differences across each dimension. This shift changes the geometry of the space, producing diamond-shaped level sets rather than circular ones, as illustrated in Figure 3. Such geometry can be advantageous in contexts involving high-dimensional data, where the Euclidean distance may become less interpretable or dominated by large outliers. The Manhattan distance also tends to preserve meaningful differences in cases where numerous features are zero-inflated, a common occurrence in microbiome data and species abundance matrices [16]. Its sensitivity to coordinate axes and additive nature sometimes makes it the preferred measure in fields like text mining and genomics, where absolute differences in counts or frequencies are more meaningful than squared deviations [16, 46].

In ecology, conservation science, and microbiome research, selecting a suitable Minkowski distance helps quantify compositional differences among communities and detect gradients of biodiversity or microbial assemblages [47]. In machine learning, the choice of $p$ influences the shape of clusters and the performance of distance-based classifiers, potentially improving outcomes when data exhibit particular distributional or structural characteristics [44].

#### 2.6 Chi-square Distance

The chi-square distance is a statistical measure that quantifies the dissimilarity between two distributions, especially in histogram comparison or compositional data. Its origins trace back to the pioneering work of British mathematician Karl Pearson in 1900 [48]. Pearson introduced the chi-square statistic as part of his development of the goodness-of-fit test, a method designed to evaluate how well observed data align with a theoretical distribution, especially within categorical data and contingency tables [48]. While his initial focus was on hypothesis testing against theoretical distribution, the chi-square statistic was later adapted as a measure for comparing empirical distributions directly, without referring to any theoretical distribution [49]. This adaptation expanded its utility across various fields, including image processing, pattern recognition, machine learning, and biological sciences, where comparing distributions of categorical data is essential [50].

The chi-square distance can be computed using the formulation from the `vegan` *R* package [51]:

$$\chi_{ij} = \sqrt{\sum_{k=1}^{K} \frac{\left(\frac{a_{ik}}{a_{i\cdot}} - \frac{a_{jk}}{a_{j\cdot}}\right)^2}{\bar{a}_{\cdot k}}} = \sqrt{\sum_{k=1}^{K} \frac{(r_{ik} - r_{jk})^2}{\bar{a}_{\cdot k}}},$$

where $\bar{a}_{\cdot k} = a_{\cdot k}/N$ and $a_{\cdot k} = \sum_i a_{ik}$. The transformation $r_{ik}/\sqrt{\bar{a}_{\cdot k}}$ is usually referred as chi-square transformation. It aggregates the values across all communities for each species $k$ and takes average according to the number of communities. This normalization term acts as a weighting factor to adjust for the relative importance of each species based on its contribution to the total abundance.

The chi-square distance has become an invaluable tool in various applications. In microbiome research, it is employed to compare the composition of microbial communities by measuring differences in the abundance of microbial taxa between samples [16]. It is also the distance function that inherent in correspondence analysis [39]. The chi-square distance enables researchers to detect significant variations in microbial populations, especially rare species that might be critical in understanding disease mechanisms or ecological dynamics [16].

#### 2.7 Hellinger Distance

Hellinger distance, established in the context of probability theory by Ernst Hellinger in 1909 [52], measures the dissimilarity between two probability distributions in a way that is both bounded and symmetric [53]. Unlike traditional measures that might magnify large discrepancies or diminish the importance of rare components, Hellinger distance applies a square-root transformation to the probability values, ensuring that both common and rare categories contribute meaningfully to the assessment. For two discrete probability distributions $P = (p_1, p_2, \ldots, p_K)$ and $Q = (q_1, q_2, \ldots, q_K)$, the Hellinger distance is defined as

$$d_{Hellinger}(P, Q) = \frac{1}{\sqrt{2}} \sqrt{\sum_{i=1}^{K} (\sqrt{p_i} - \sqrt{q_i})^2}.$$

This formulation constrains the distance to lie within the interval $[0, 1]$, with 0 indicating identical distributions and 1 representing maximum divergence.

Hellinger distance has found broad utility in ecology, microbiome research, and other biological and environmental sciences [54], where the underlying data often represent relative abundances rather than absolute counts,

$$d_{Hellinger}(i, j) = \frac{1}{\sqrt{2}} \sqrt{\sum_{k=1}^{K} (\sqrt{r_{ik}} - \sqrt{r_{jk}})^2}.$$

By incorporating a square-root transformation, the measure downweights dominant taxa and alleviates the influence of very small proportions, improving robustness against sampling errors and compositional biases. In statistical inference and machine learning, it provides stable and interpretable comparisons between distributions, facilitates clustering and classification tasks, and can enhance performance in non-parametric estimation [55–57]. Its versatile nature and sound mathematical properties make Hellinger distance a valuable tool for capturing patterns and differences in complex, probability-based data sets [58].

#### 2.8 Mahalanobis Distance

Mahalanobis distance, introduced by Prasanta Chandra Mahalanobis in 1936 [59], provides a multivariate measure of dissimilarity that accounts for correlations among variables. Unlike Euclidean distance, which treats each dimension independently, Mahalanobis distance adjusts for the covariance structure of the variables, giving more balanced, scale-invariant assessments of how similar or different points are in a multidimensional space. For two vectors $x = (x_1, x_2, \ldots, x_K)$ and $y = (y_1, y_2, \ldots, y_K)$, and an variance–covariance matrix $\Sigma$ [60], the Mahalanobis distance is defined as

$$d_M(x, y) = \sqrt{(x - y)^T \Sigma^{-1} (x - y)}.$$

This formulation not only re-scales features according to their variances, but also reduces redundancy by down-weighting directions of high correlation. Its use spans fields such as ecology, where it helps discern ecological niches by comparing species assemblages under correlated environmental gradients [16], and in pattern recognition, machine learning, and outlier detection, where capturing the “shape” and internal structure of data is critical. Mahalanobis distance’s ability to incorporate covariance information makes it a powerful tool for multivariate analyses, guiding the identification of clusters, discriminating among groups, and improving classification accuracy in complex, high-dimensional data sets.

In microbiome analyses, each dimension commonly represents an OTU (Operational Taxonomic Unit), and the matrix $\Sigma$ reflects the variance and covariance of these OTUs across samples,

$$d_M(i,j) = \sqrt{\sum_{k=1}^{K}\sum_{l=1}^{K}(a_{ik}-a_{jk})(\Sigma^{-1})_{k,l}(a_{il}-a_{jl})},$$

where $(\Sigma^{-1})_{k,l}$ is the $(k,l)$-th element of $\Sigma^{-1}$. Notably, as R packages vegan [61] and phyloseq [62] do, we replace the true covariance matrix $\Sigma$ with the sample covariance matrix $S$. It is derived from the abundance patterns of OTUs rather than their representative sequences, while Pavoine et al. 2004 [63] interprets $\Sigma_{k,l}$ as the distance between species $k$ and $l$. In microbial applications, since number of OTUs is usually larger than number of samples, the inverse of $\Sigma$ does not exist. Alternatively, general inverse could be considered. By incorporating these abundance-based covariance structures, Mahalanobis distance can capture the correlated nature of microbial communities more accurately. Literature on multivariate analyses in microbial ecology [16, 64] underscores the importance of considering abundance-based covariance patterns, as they help in revealing meaningful ecological interactions and community-level structures.

#### 2.9 Aitchinson Distance

Aitchison distance, introduced by John Aitchison in 1982 [65] in the context of compositional data analysis, provides a meaningful notion of distance that respects the simplex structure inherent to compositional vectors. Traditional distances defined on the Euclidean space, such as the Minkowski distances, are not well-suited for compositional data. This is because compositional data are characterized by nonnegative components that sum to a constant. Moreover, the unit-sum constraint and differences in absolute scale can further distort the distances when these traditional measures are applied. The Aitchison distance circumvents these issues by mapping compositions into a log-ratio space, where standard Euclidean geometry can be applied. The Aitchison distance is defined as

$$d_A(i,j) = \left(\sum_{k=1}^{K}\left(\log\frac{r_{ik}}{g(i)} - \log\frac{r_{jk}}{g(j)}\right)^2\right)^{\frac{1}{2}},$$

where

$$g(i) = \left(\prod_{k=1}^{K} r_{ik}\right)^{\frac{1}{K}}.$$

This formulation treats ratios of parts, rather than their absolute differences, ensuring that the analysis focuses on relative information—crucial in fields such as geochemistry or microbiome research, where relative abundances carry more interpretative significance than absolute counts [66, 67].

In practice, compositional data are often characterized by the presence of zeros, the occurrence of outliers, and heavy-tailed distributions. To address such complexities, robust versions of the Aitchison distance have been developed [68]. These variants incorporate robust estimators of center and scatter, or employ zero-replacement techniques that maintain the compositional nature of the data while diminishing the influence of problematic values. By reducing sensitivity to extreme observations and properly handling zeros, robust Aitchison distances enhance stability and reliability in real-world applications, improving interpretations of ecological niches, dietary patterns, environmental pollution gradients, and microbial community structures.

### 3 Construction of Dissimilarity Measures

| Category | Space | Class | Dissimilarity | Metric? | Euclidean? |
|---|---|---|---|---|---|
| scale difference | $\mathbb{R}^n$ or $S^n$ | $D_1(\mathbf{x},\mathbf{y}) = \lVert\psi_1(\mathbf{x}) - \psi_1(\mathbf{y})\rVert_p$ | Euclidean | Yes | Yes |
| | | | Manhattan | Yes | No |
| | | | Mahalanobis | Yes | Yes |
| | | | Chord | Yes | Yes |
| | | | Aitchison | Yes | Yes |
| | | | Robust Aitchison | Yes | Yes |
| | | | Chi-square | Yes | Yes |
| difference scale | $\mathbb{R}^n$ or $S^n$ | $D_2(\mathbf{x},\mathbf{y}) = \lVert\psi_2(\mathbf{x}-\mathbf{y})\rVert_p$ | Bray-Curtis | No | No |
| | | | Kulczyński | Unclear | Unclear |
| | | | Weighted UniFrac | Likely | No |
| presence difference | $\mathbb{H}^n$ | $D_3(\mathbf{x},\mathbf{y}) = 1 - \frac{\mathbf{x}\cdot\mathbf{y}}{\psi_H(\mathbf{x},\mathbf{y})}$ | Jaccard | Yes | No |
| | | | Unweighted UniFrac | Yes | No |
| distribution difference | Distributions | $D_f(P\Vert Q) = \int_{\mathcal{X}} f\left(\frac{dP}{dQ}\right) dQ$ | Hellinger | Yes | Yes |
| | | | Jensen-Shannon | Yes | Unclear |

**Table 1** Classification of commonly used dissimilarity measures. Measures Category 1 operate in Euclidean space (abundance data) or compositional simplex (relative abundance data) and compute the $L_p$ norm after applying a normalization function $\psi_1$ to each data point individually before calculating the difference. Measures in Category 2 also operate in Euclidean space or compositional simplex but differ by first computing the difference between data points and then applying a normalization function $\psi_2$. Measures in Category 3 are Defined in the Hamming space $\mathbb{H}^n$ and usually assess dissimilarity based on presence-absence data. Measures in Category 4 measures the dissimilarity between probability mass functions, which can also be applied to relative abundance data.

Table 1 presents a classification of commonly used dissimilarity measures. In this section, we present the detailed constructions of each dissimilarity measures.

The first category consists of dissimilarity measures that normalize the abundance by normalization function $\psi_1(\cdot)$ or relative abundance vectors separately and applies $L_p$ norm to evaluate there difference. We list the corresponding normalization functions of the dissimilarity measures as follows:

- Euclidean, $\psi_1(\mathbf{x}) = \mathbf{x}$, $p = 2$.
- Manhattan, $\psi_1(\mathbf{x}) = \mathbf{x}$, $p = 1$.
- Mahalanobis, $\psi_1(\mathbf{x}) = \Sigma^{-1/2}\mathbf{x}$, $p = 2$.

- Chord, $\psi_1(\mathbf{x}) = \mathbf{x}/\left\|\mathbf{x}\right\|_2$, $p = 2$.
- Aitchison, $\psi_1(\mathbf{x}) = \text{CLR}(\mathbf{x})$, $p = 2$, $\mathbf{x} \in S^n$, where CLR($\mathbf{x}$) represents the CLR transformation.
- Robust Aitchison, $\psi_1(\mathbf{x}) = \Sigma^{-1/2}\text{CLR}(\mathbf{x})$, $\mathbf{x} \in S^n$.
- Chi-square, $\psi_1(\mathbf{x}) = \chi(\mathbf{x})$, $p = 2$, $\mathbf{x} \in S^n$, where $\chi(\mathbf{x})$ represents the chi-square transformation.

The second category of dissimilarity measures normalizes the elementwise differences in relative abundances before applying the $L_p$ norm. We list the detailed construction of dissimilarity measures in category 2 as follows:

- Bray-Curtis, $\psi_2(\mathbf{x} - \mathbf{y}) = |\mathbf{x} - \mathbf{y}|/\left\|\mathbf{x} + \mathbf{y}\right\|_1$, $p = 1$.
- Kulczyński, $\psi_2(\mathbf{x} - \mathbf{y}) = |\mathbf{x} - \mathbf{y}|/\frac{2}{\|\mathbf{x}\|_1 + \|\mathbf{y}\|_1}$, $p = 1$.
- Weighted UniFrac, $\psi_2(\mathbf{x} - \mathbf{y}) = \mathbf{b} \circ |\mathbf{x} - \mathbf{y}|/\left\|\mathbf{b} \circ (\mathbf{x} + \mathbf{y})\right\|_1$, $p = 1$ where $\mathbf{b}$ represents the vector of branch length from each nodes to their nearest ancestor nodes and $\circ$ represents Hadamard product (element-wise product).

The key distinction between these two categories lies in whether normalization is applied directly to the raw data before computing differences or to the differences of the raw data themselves. Hence, the theoretical justification of the dissimilarity in the second category is difficult. If we use relative abundance, the denominator (total abundance) in the Bray–Curtis formula becomes constant (usually 2), simplifying the math:

$$\text{BC}_{ij} = \frac{\sum_k (r_{ik} - r_{jk})}{2},$$
$$\text{BC}_{ij} = 1 - \sum_k \min(r_{ik}, r_{jk}),$$

which becomes a scaled Manhattan distance. Bray–Curtis on relative abundances may reduce the impact of sample depth differences, which is good when you want to compare community structure rather than raw load. Compositionality still matters, since relative abundances are constrained (they must sum to 1). Some researchers prefer log-ratio transformations (e.g., CLR) for downstream analyses like clustering or regression.

Category 3 includes dissimilarity measures defined in Hamming space and is usually applied to presence-absence data. Since the Hamming inner product $\mathbf{x} \cdot \mathbf{y}$ is usually used as a measure of similarity of two Hamming vectors, the dissimilarity measures in this category is one minus the normalized similarity. For Jaccard, the normalization function is $\psi_H(\mathbf{x}, \mathbf{y}) = \sum_i I(x_i + y_i > 0)$. For unweighted UniFrac, since there are branch length assigned to each OTU, it does not follow the form listed in the table. Instead, it requires a little modification

$$\begin{aligned} d_{\text{Unweighted UniFrac}}(\mathbf{x}, \mathbf{y}) &= \frac{\sum_i b_k[I(x_i + y_i > 0) - x_i y_i]}{\sum_i b_i I(x_i + y_i > 0)} \\ &= 1 - \frac{\sum_i b_k x_i y_i}{\sum_i b_i I(x_i + y_i > 0)}, \end{aligned}$$

which makes it similar to a weighted version of Jaccard. The theoretical justification of dissimilarity measures is also difficult. Though there are some proof existing, most of them are still not Euclidean due to the data in Hamming space having different geometrical feature compared to Euclidean space. Hence, diagnostics is also suggested before applying.

The fourth category takes use of the concept of $f$-divergence and use relative abundances as probability mass function. In its mathematical formulation, the Radon-Nikodym derivative [69] naturally works as a measure of dissimilarity between two probability measures. Hence, we regard the dissimilarity measures in the fourth category as average normalized measures based on Radon-Nikodym derivative, where the average is weighted average based on probability measure $Q$ and $f$ works as the normalization function. When applied to relative abundance data, the general form of the fourth category is

$$D_f(i,j) = \sum_{k=1}^{K} r_{jk} f(\frac{r_{ik}}{r_{jk}}).$$

However, not all dissimilarity measures under this category are metric. Theoretical justification or diagnostics are required before applying them to downstream analysis.

### 4 Supplementary Results: Metric Property and Euclidean Deviation (FNI) of Dissimilarity Measures

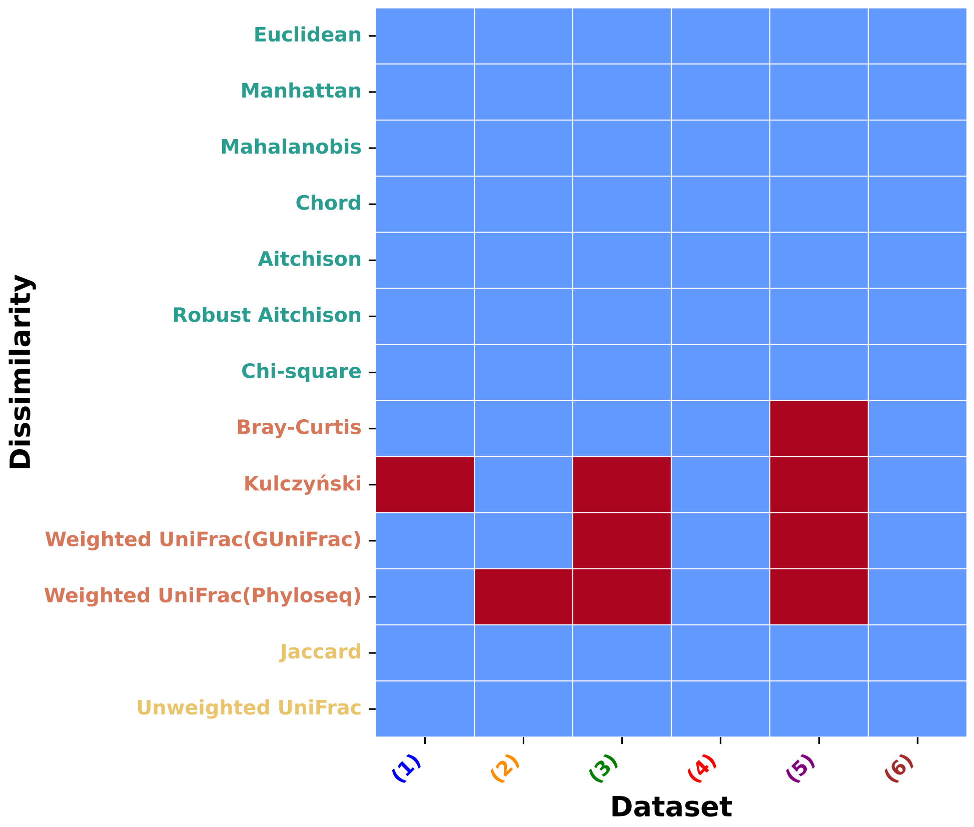


**Fig. 4** *Metric property of dissimilarity measures across datasets.* This heatmap indicates whether each dissimilarity measure satisfies the metric property within each dataset. Blue squares denote that the dissimilarity is metric; red squares indicate violations of the metric condition. Dissimilarities are grouped and color-coded by category to facilitate interpretation across types of measures.

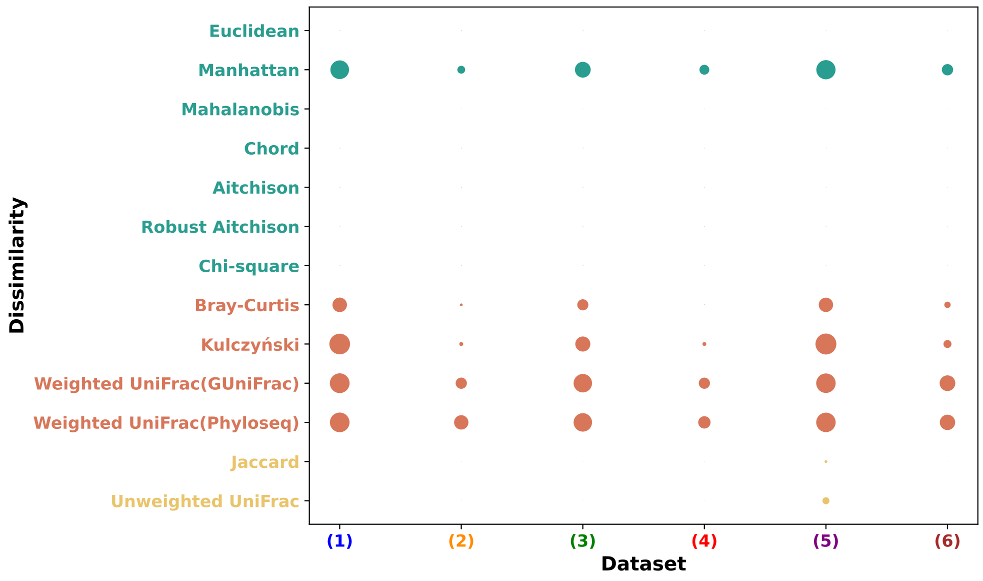


**Fig. 5** *Euclidean deviation (FNI) of dissimilarity measures across datasets.* The bubble plot visualizes the Fraction of Negative inertia (FNI) for each dissimilarity measure across datasets. Larger bubbles correspond to higher FNI values, indicating greater deviation from Euclidean geometry. An FNI of zero implies that the dissimilarity is Euclidean for that dataset. Dissimilarity labels are color-coded by category.

#### 5 Diagnostics and Remedy Results

In summary, the diagnostics across datasets (Tables 2, 3, 4, 5, 6, 7) show that classic metric choices—Manhattan, Euclidean, chord, Hellinger, (robust) Aitchison, Jaccard, binomial, CAO, and $\chi^2$—are consistently flagged as metrics and yield zero PCoA negative inertia (Euclidean invariably at 0%), whereas several abundance-weighted or alternative indices (e.g., altGower, Morisita, Horn, and occasionally Kulczyński) are non-metric and exhibit sizeable negative inertia fractions (often $\sim$15–21%); Chao and Raup are among the most problematic, with large NegCounter/ZeroCounter values and the highest negative fractions (up to $\sim$48–49%). Without remedies (Table 8), non-phylogenetic performers are typically Bray–Curtis and CAO, while phylogenetic measures—especially weighted UniFrac—can be strongest when the phylogeny aligns with signal (e.g., Pig Gut: weighted UniFrac pseudo-$F! =!10.92$, pseudo-$R^2 = 0.2007$, MiRKAT $R^2 = 0.1895$), and this pattern broadly persists across additional datasets (including IBD_wgs). Applying the Higham projection (Table 9) modestly increases or stabilizes pseudo-$F$/pseudo-$R^2$ for many distances (notably boosting phylogenetic measures in Covid and Pig Gut; e.g., Pig Gut weighted UniFrac to pseudo-$F = 11.67$, pseudo-$R^2 = 0.2116$), while MiRKAT $R^2$ tracks these gains. In contrast, Tikhonov regularization (Table 10) generally shrinks effect sizes—most visibly for weighted UniFrac (e.g., Pig Gut pseudo-$F$ drops to 4.93, pseudo-$R^2$ to 0.1017)—yet largely preserves the within-dataset ranking (Bray/CAO among top non-phylogenetic; weighted UniFrac often leading when appropriate). Together, these results indicate: (i) metric validity and Euclidean embeddability issues are concentrated in a subset of abundance-weighted/coverage-based indices, (ii) Higham correction is preferable when maintaining or slightly enhancing signal while enforcing PSD structure, and (iii) Tikhonov offers a conservative, stabilizing alternative at the cost of attenuated test statistics.

| Dissimilarity | IsMetric | NegCounter | ZeroCounter | PcoaNegFrac | pseudo_F | pseudo_R2 | p_value | MiRKAT_Rsquared |
|---|---|---|---|---|---|---|---|---|
| manhattan | YES | 0 | 0 | 12.06% | 1.31023 | 0.05736 | 0.03203 | 0.00301 |
| euclidean | YES | 0 | 0 | 0% | 1.97807 | 0.08413 | 0.003 | 0.00366 |
| canberra | YES | 0 | 0 | 0% | 1.44541 | 0.0629 | 0 | 0.00303 |
| clark | YES | 0 | 0 | 0% | 1.29833 | 0.05687 | 0 | 0.00303 |
| bray | YES | 0 | 0 | 7.31% | 1.61735 | 0.06986 | 0 | 0.003 |
| kulczynski | NO | 0 | 0 | 14.82% | 1.36959 | 0.0598 | 0 | 0.0025 |
| jaccard | YES | 0 | 0 | 0.00% | 1.51568 | 0.06576 | 0 | 0.0033 |
| gower | YES | 0 | 0 | 3.42% | 1.19847 | 0.05272 | 0.08909 | 0.00213 |
| altGower | NO | 0.0042 | 0 | 18.29% | 1.35481 | 0.05919 | 0.03804 | 0.00393 |
| morisita | NO | 0.00381 | 0 | 19.14% | 1.33773 | 0.05849 | 0.02803 | 0.00183 |
| horn | NO | 0.0038 | 0 | 19.14% | 1.33774 | 0.05849 | 0.03303 | 0.00183 |
| binomial | YES | 0 | 0 | 0.67% | 1.76927 | 0.07593 | 0 | 0.00261 |
| chao | NO | 0.27794 | 0.00085 | 47.81% | 0.65216 | 0.0294 | 0.58458 | 0.00052 |
| cao | YES | 0 | 0 | 0.77% | 2.01599 | 0.08561 | 0 | 0.00409 |
| chisq | YES | 0 | 0 | 0% | 1.1762 | 0.05179 | 0.04304 | 0.00261 |
| chord | YES | 0 | 0 | 0% | 1.58413 | 0.06853 | 0 | 0.00309 |
| hellinger | YES | 0 | 0 | 0% | 1.81249 | 0.07764 | 0 | 0.0037 |
| aitchison | YES | 0 | 0 | 0% | 1.81249 | 0.07764 | 0 | 0.0037 |
| robust.aitchison | YES | 0 | 0 | 0% | 1.61899 | 0.06993 | 0 | 0.00324 |
| unifrac_unweighted | YES | 0 | 0 | 0% | 1.68363 | 0.07252 | 0 | 0.00307 |
| unifrac_weighted_phyloseq | YES | 0 | 0 | 13.38% | 1.44689 | 0.06296 | 0 | 0.00226 |
| unifrac_weighted_gunifrac | YES | 0 | 0 | 13.52% | 1.7549 | 0.07536 | 0.001 | 0.00353 |

**Table 2** Diagnostic results for each dissimilarity measure applied to the cardiovascular dataset.

| Dissimilarity | IsMetric | NegCounter | ZeroCounter | PcoaNegFrac | pseudo_F | pseudo_R2 | p_value | MiRKAT_Rsquared |
|---|---|---|---|---|---|---|---|---|
| manhattan | YES | 0 | 0 | 12.78% | 0.83854 | 0.042 | 0.81481 | 0.00298 |
| euclidean | YES | 0 | 0 | 0% | 0.718 | 0.03618 | 0.87087 | 0.00276 |
| canberra | YES | 0 | 0 | 0% | 1.10292 | 0.05452 | 0.00901 | 0.00661 |
| clark | YES | 0 | 0 | 0% | 1.07328 | 0.05314 | 0.01401 | 0.0065 |
| bray | NO | 0 | 0 | 7.16% | 0.93848 | 0.04678 | 0.61361 | 0.0047 |
| kulczynski | NO | 2E-05 | 0 | 15.17% | 0.86319 | 0.04318 | 0.81281 | 0.00287 |
| jaccard | YES | 0 | 0 | 0.27% | 1.00561 | 0.04995 | 0.44344 | 0.00595 |
| gower | YES | 0 | 0 | 6.25% | 0.87476 | 0.04374 | 0.8038 | 0.00289 |
| altGower | NO | 0.01778 | 0 | 21.43% | 0.79817 | 0.04006 | 0.76877 | 8E-04 |
| morisita | NO | 0.05391 | 1E-05 | 18.56% | 0.73842 | 0.03718 | 0.81882 | 0.00311 |
| horn | NO | 0.05233 | 0 | 18.14% | 0.74234 | 0.03736 | 0.82883 | 0.00314 |
| mountford | YES | 0 | 0 | 0% | 1.00767 | 0.05005 | 0.02503 | 0.00623 |
| raup | NO | 0.05594 | 0.81819 | 43.93% | 0.47945 | 0.02446 | 0.58759 | -3E-05 |
| binomial | YES | 0 | 0 | 1.42% | 1.06768 | 0.05287 | 0.24625 | 0.00608 |
| chao | NO | 0.02922 | 0 | 28.05% | 0.82996 | 0.04159 | 0.69369 | 0.00365 |
| cao | YES | 0 | 0 | 0.83% | 1.14492 | 0.05648 | 0.11512 | 0.00556 |
| mahalanobis | YES | 0 | 0 | 0% | 1.00277 | 0.04982 | 0.4975 | 0.0061 |
| chisq | YES | 0 | 0 | 0% | 0.60893 | 0.03086 | 0.88088 | 0.00479 |
| chord | YES | 0 | 0 | 0% | 0.86241 | 0.04315 | 0.75475 | 0.00572 |
| hellinger | YES | 0 | 0 | 0% | 1.07461 | 0.0532 | 0.25425 | 0.00694 |
| aitchison | YES | 0 | 0 | 0% | 1.07461 | 0.0532 | 0.24725 | 0.00694 |
| robust.aitchison | YES | 0 | 0 | 0% | 1.07044 | 0.053 | 0.14014 | 0.00622 |
| unifrac_unweighted | YES | 0 | 0 | 1.70% | 1.22786 | 0.06033 | 0.1011 | 0.00659 |
| unifrac_weighted_phyloseq | NO | 2E-05 | 0 | 13.15% | 0.86975 | 0.0435 | 0.68569 | 0.00283 |
| unifrac_weighted_gunifrac | NO | 2E-05 | 0 | 13.15% | 0.86975 | 0.0435 | 0.65566 | 0.00283 |

**Table 3** Diagnostic results for each dissimilarity measure applied to the Concordance oral 16S dataset.

| Dissimilarity | IsMetric | NegCounter | ZeroCounter | PcoaNegFrac | pseudo_F | pseudo_R2 | p_value | MiRKAT_Rsquared |
|---|---|---|---|---|---|---|---|---|
| manhattan | YES | 0 | 0 | 2.14% | 1.47468 | 0.16563 | 0.001 | 0.01333 |
| euclidean | YES | 0 | 0 | 0% | 1.21405 | 0.14047 | 0.04204 | 0.0174 |
| canberra | YES | 0 | 0 | 0% | 1.15832 | 0.13489 | 0 | 0.01619 |
| clark | YES | 0 | 0 | 0% | 1.10741 | 0.12973 | 0 | 0.01639 |
| bray | YES | 0 | 0 | 0.26% | 1.4536 | 0.16365 | 0 | 0.01452 |
| kulczynski | YES | 0 | 0 | 0.54% | 1.42815 | 0.16125 | 0 | 0.01372 |
| jaccard | YES | 0 | 0 | 0% | 1.29266 | 0.14822 | 0 | 0.01529 |
| gower | YES | 0 | 0 | 4.32% | 1.09522 | 0.12849 | 0.08008 | 0.01583 |
| altGower | NO | 0.01147 | 0 | 20.84% | 0.88974 | 0.10696 | 0.66667 | 0.01218 |
| morisita | NO | 0.00039 | 0 | 8.23% | 1.20259 | 0.13933 | 0.07808 | 0.01109 |
| horn | NO | 0.00036 | 0 | 8.20% | 1.20282 | 0.13935 | 0.08809 | 0.0111 |
| mountford | YES | 0 | 0 | 0% | 1.00492 | 0.11916 | 0 | 0.01692 |
| raup | NO | 0.11836 | 0.64476 | 49.15% | 0.41495 | 0.0529 | 0.13714 | -0.00886 |
| binomial | YES | 0 | 0 | 0.81% | 1.48165 | 0.16629 | 0 | 0.01586 |
| chao | NO | 0.00176 | 0 | 17.15% | 1.43091 | 0.16151 | 0.00801 | 0.01014 |
| cao | YES | 0 | 0 | 0% | 1.46083 | 0.16433 | 0 | 0.01444 |
| mahalanobis | YES | 0 | 0 | 0% | 1 | 0.11864 | 0.01502 | 0.01695 |
| chisq | YES | 0 | 0 | 0% | 1.25247 | 0.14428 | 0 | 0.01826 |
| chord | YES | 0 | 0 | 0% | 1.20488 | 0.13956 | 0.02302 | 0.01505 |
| hellinger | YES | 0 | 0 | 0% | 1.4194 | 0.16042 | 0 | 0.01354 |
| aitchison | YES | 0 | 0 | 0% | 1.57558 | 0.17498 | 0 | 0.01348 |
| robust.aitchison | YES | 0 | 0 | 0% | 1.31388 | 0.15029 | 0 | 0.01506 |
| unifrac_unweighted | YES | 0 | 0 | 0% | 1.30058 | 0.14899 | 0 | 0.01507 |
| unifrac_weighted_phyloseq | NO | 1E-04 | 0 | 7.14% | 1.85404 | 0.19973 | 0.001 | 0.00882 |
| unifrac_weighted_gunifrac | YES | 0 | 0 | 4.38% | 1.2198 | 0.14104 | 0.13013 | 0.01377 |

**Table 4** Diagnostic results for each dissimilarity measure applied to the Covid dataset.

| Dissimilarity | IsMetric | NegCounter | ZeroCounter | PcoaNegFrac | pseudo_F | pseudo_R2 | p_value | MiRKAT_Rsquared |
|---|---|---|---|---|---|---|---|---|
| manhattan | YES | 0 | 0 | 4.41% | 1.01723 | 0.07647 | 0.43043 | 0.01464 |
| euclidean | YES | 0 | 0 | 0% | 0.93658 | 0.07083 | 0.55455 | 0.01338 |
| canberra | YES | 0 | 0 | 0% | 1.1111 | 0.08294 | 0 | 0.01287 |
| clark | YES | 0 | 0 | 0% | 1.07363 | 0.08037 | 0 | 0.01215 |
| bray | YES | 0 | 0 | 1.41% | 1.2506 | 0.09239 | 0.02603 | 0.01463 |
| kulczynski | YES | 0 | 0 | 2.23% | 1.25109 | 0.09242 | 0.02102 | 0.01513 |
| jaccard | YES | 0 | 0 | 0% | 1.17051 | 0.08699 | 0.01201 | 0.01397 |
| gower | YES | 0 | 0 | 8.16% | 1.06061 | 0.07947 | 0.35936 | 0.01249 |
| altGower | NO | 0.013 | 0 | 17.59% | 0.74262 | 0.057 | 0.83483 | 0.00394 |
| morisita | NO | 0.00208 | 0 | 12.99% | 1.1499 | 0.08559 | 0.19119 | 0.01176 |
| horn | NO | 0.00205 | 0 | 12.97% | 1.15015 | 0.0856 | 0.21722 | 0.01176 |
| mountford | YES | 0 | 0 | 0% | 1.00371 | 0.07553 | 0.001 | 0.01081 |
| binomial | YES | 0 | 0 | 2.55% | 1.18799 | 0.08817 | 0.11912 | 0.01599 |
| chao | NO | 0.00069 | 0 | 20.43% | 1.02449 | 0.07697 | 0.41742 | 0.01097 |
| cao | YES | 0 | 0 | 1.31% | 1.17225 | 0.0871 | 0.01902 | 0.01548 |
| mahalanobis | YES | 0 | 0 | 0% | 1 | 0.07527 | 0.86486 | 0.01075 |
| chisq | YES | 0 | 0 | 0% | 1.12248 | 0.08372 | 0.1962 | 0.01531 |
| chord | YES | 0 | 0 | 0% | 1.26863 | 0.0936 | 0.03403 | 0.01555 |
| hellinger | YES | 0 | 0 | 0% | 1.22859 | 0.09091 | 0.01201 | 0.01417 |
| aitchison | YES | 0 | 0 | 0% | 1.22859 | 0.09091 | 0.02102 | 0.01417 |
| robust.aitchison | YES | 0 | 0 | 0% | 1.09049 | 0.08152 | 0.0961 | 0.01285 |
| unifrac_unweighted | YES | 0 | 0 | 0% | 1.16122 | 0.08636 | 0.003 | 0.01401 |
| unifrac_weighted_phyloseq | YES | 0 | 0 | 8.19% | 0.93881 | 0.07099 | 0.49249 | 0.00786 |
| unifrac_weighted_gunifrac | YES | 0 | 0 | 8.52% | 1.02873 | 0.07726 | 0.42843 | 0.00957 |

**Table 5** Diagnostic results for each dissimilarity measure applied to the HIV Gut dataset.

| Dissimilarity | IsMetric | NegCounter | ZeroCounter | PcoaNegFrac | pseudo_F | pseudo_R2 | p_value | MiRKAT_Rsquared |
|---|---|---|---|---|---|---|---|---|
| manhattan | YES | 0 | 0 | 8.61% | 1.28003 | 0.09639 | 0.02603 | 0.01228 |
| euclidean | YES | 0 | 0 | 0% | 1.38656 | 0.10358 | 0.06907 | 0.02246 |
| canberra | YES | 0 | 0 | 0% | 1.13487 | 0.0864 | 0 | 0.00663 |
| clark | YES | 0 | 0 | 0% | 1.09342 | 0.08351 | 0 | 0.0063 |
| bray | YES | 0 | 0 | 4.27% | 1.38867 | 0.10372 | 0 | 0.01314 |
| kulczynski | NO | 0 | 0 | 7.85% | 1.34608 | 0.10086 | 0 | 0.0116 |
| jaccard | YES | 0 | 0 | 0% | 1.32151 | 0.0992 | 0 | 0.01108 |
| gower | NO | 0 | 0 | 10.53% | 0.76898 | 0.06022 | 0.91091 | 0.00336 |
| altGower | NO | 0.04493 | 0 | 38.79% | 0.76914 | 0.06023 | 0.66867 | -0.01276 |
| morisita | NO | 0.00249 | 0 | 13.81% | 1.24427 | 0.09395 | 0.01802 | 0.0157 |
| horn | NO | 0.00244 | 0 | 13.55% | 1.24715 | 0.09414 | 0.01201 | 0.01583 |
| mountford | NO | 0.00024 | 0 | 0.42% | 1.02928 | 0.079 | 0.001 | 0.00551 |
| raup | NO | 0.24643 | 0.25558 | 48.09% | 1.01741 | 0.07816 | 0.50851 | -0.00176 |
| binomial | YES | 0 | 0 | 3.67% | 0.97348 | 0.07504 | 0.50751 | 0.00545 |
| chao | NO | 0.00327 | 0 | 25.10% | 1.09752 | 0.0838 | 0.17718 | 0.00996 |
| cao | NO | 0 | 0 | 2.40% | 1.32689 | 0.09956 | 0 | 0.00889 |
| mahalanobis | YES | 0 | 0 | 0% | 0.99124 | 0.0763 | 1 | 0.00549 |
| chisq | YES | 0 | 0 | 0% | 0.58018 | 0.04612 | 0.63063 | 0.00634 |
| chord | YES | 0 | 0 | 0% | 1.37654 | 0.10291 | 0 | 0.01682 |
| hellinger | YES | 0 | 0 | 0% | 1.45619 | 0.10822 | 0 | 0.01342 |
| aitchison | YES | 0 | 0 | 0% | 1.45619 | 0.10822 | 0 | 0.01342 |
| robust.aitchison | YES | 0 | 0 | 0% | 1.13335 | 0.0863 | 0.02002 | 0.00676 |
| unifrac_unweighted | YES | 0 | 0 | 0% | 1.197 | 0.0907 | 0 | 0.00899 |
| unifrac_weighted_phyloseq | NO | 0.00209 | 0 | 11.81% | 0.98643 | 0.07596 | 0.50951 | 0.01034 |
| unifrac_weighted_gunifrac | NO | 0 | 0 | 11.80% | 1.4507 | 0.10785 | 0.002 | 0.01908 |

**Table 6** Diagnostic results for each dissimilarity measure applied to the IBD 16S dataset.

| Dissimilarity | IsMetric | NegCounter | ZeroCounter | PcoaNegFrac | pseudo_F | pseudo_R2 | p_value | MiRKAT_Rsquared |
|---|---|---|---|---|---|---|---|---|
| manhattan | YES | 0 | 0 | 3.38% | 4.05871 | 0.08534 | 0 | 0.0651 |
| euclidean | YES | 0 | 0 | 0% | 3.49776 | 0.07442 | 0 | 0.05306 |
| canberra | YES | 0 | 0 | 0% | 2.45593 | 0.05344 | 0 | 0.03723 |
| clark | YES | 0 | 0 | 0% | 1.95491 | 0.04301 | 0 | 0.02845 |
| bray | YES | 0 | 0 | 0.02% | 5.46593 | 0.11163 | 0 | 0.09123 |
| kulczynski | YES | 0 | 0 | 0.52% | 5.76707 | 0.11706 | 0 | 0.09717 |
| jaccard | YES | 0 | 0 | 0% | 3.54906 | 0.07543 | 0 | 0.05892 |
| gower | YES | 0 | 0 | 2.68% | 2.76856 | 0.05984 | 0 | 0.04159 |
| altGower | NO | 0.00085 | 0 | 9.30% | 1.94848 | 0.04287 | 0 | 0.00118 |
| morisita | NO | 2E-04 | 0 | 9.48% | 7.39691 | 0.14533 | 0 | 0.12498 |
| horn | NO | 0.00019 | 0 | 9.41% | 7.38988 | 0.14521 | 0 | 0.12484 |
| mountford | YES | 0 | 0 | 0% | 1.01201 | 0.02274 | 0 | 0.01145 |
| binomial | YES | 0 | 0 | 0% | 4.65826 | 0.09673 | 0 | 0.07457 |
| chao | NO | 2E-05 | 0 | 11.54% | 4.89073 | 0.10107 | 0 | 0.08074 |
| cao | YES | 0 | 0 | 0% | 4.19175 | 0.08789 | 0 | 0.06694 |
| mahalanobis | YES | 0 | 0 | 0% | 1 | 0.02247 | 0.81281 | 0.01124 |
| chisq | YES | 0 | 0 | 0% | 2.34627 | 0.05118 | 0 | 0.03577 |
| chord | YES | 0 | 0 | 0% | 5.6534 | 0.11502 | 0 | 0.09498 |
| hellinger | YES | 0 | 0 | 0% | 5.10108 | 0.10496 | 0 | 0.086 |
| aitchison | YES | 0 | 0 | 0% | 5.10108 | 0.10496 | 0 | 0.086 |
| robust.aitchison | YES | 0 | 0 | 0% | 3.32876 | 0.07108 | 0 | 0.05584 |
| unifrac_unweighted | YES | 0 | 0 | 0% | 3.37326 | 0.07197 | 0 | 0.05326 |
| unifrac_weighted_phyloseq | YES | 0 | 0 | 5.27% | 10.92424 | 0.20072 | 0 | 0.18948 |
| unifrac_weighted_gunifrac | YES | 0 | 0 | 4.34% | 8.93685 | 0.17043 | 0 | 0.14691 |

**Table 7** Diagnostic results for each dissimilarity measure applied to the Pig Gut dataset.

| | Dissimilarity | pseudo_F | pseudo_R2 | MiRKAT_Rsquared |
|---|---|---|---|---|
| cardiovascular | bray | 1.61735 | 0.06986 | 0.003 |
| Concordance_oral_16S | bray | 0.93848 | 0.04678 | 0.0047 |
| covid | bray | 1.4536 | 0.16365 | 0.01452 |
| HIV_gut | bray | 1.2506 | 0.09239 | 0.01463 |
| IBD_16s | bray | 1.38867 | 0.10372 | 0.01314 |
| IBD_wgs | bray | 1.31194 | 0.10542 | 0.01291 |
| Pig_gut | bray | 5.46593 | 0.11163 | 0.09123 |
| cardiovascular | cao | 2.01599 | 0.08561 | 0.00409 |
| Concordance_oral_16S | cao | 1.14492 | 0.05648 | 0.00556 |
| HIV_gut | cao | 1.17225 | 0.0871 | 0.01548 |
| IBD_16s | cao | 1.32689 | 0.09956 | 0.00889 |
| IBD_wgs | cao | 1.27089 | 0.10246 | 0.01422 |
| cardiovascular | chao | 0.65216 | 0.0294 | 0.00052 |
| Concordance_oral_16S | chao | 0.82996 | 0.04159 | 0.00365 |
| covid | chao | 1.43091 | 0.16151 | 0.01014 |
| HIV_gut | chao | 1.02449 | 0.07697 | 0.01097 |
| IBD_16s | chao | 1.09752 | 0.0838 | 0.00996 |
| IBD_wgs | chao | 0.75732 | 0.06369 | 0.0096 |
| Pig_gut | chao | 4.89073 | 0.10107 | 0.08074 |
| cardiovascular | jaccard | 1.51568 | 0.06576 | 0.0033 |
| Concordance_oral_16S | jaccard | 1.00561 | 0.04995 | 0.00595 |
| Concordance_oral_16S | unifrac_unweighted | 1.22786 | 0.06033 | 0.00659 |
| cardiovascular | unifrac_weighted_gunifrac | 1.7549 | 0.07536 | 0.00353 |
| Concordance_oral_16S | unifrac_weighted_gunifrac | 0.86975 | 0.0435 | 0.00283 |
| covid | unifrac_weighted_gunifrac | 1.2198 | 0.14104 | 0.01377 |
| HIV_gut | unifrac_weighted_gunifrac | 1.02873 | 0.07726 | 0.00957 |
| IBD_16s | unifrac_weighted_gunifrac | 1.4507 | 0.10785 | 0.01908 |
| IBD_wgs | unifrac_weighted_gunifrac | 1.30498 | 0.10492 | 0.02381 |
| Pig_gut | unifrac_weighted_gunifrac | 8.93685 | 0.17043 | 0.14691 |
| cardiovascular | unifrac_weighted_phyloseq | 1.44689 | 0.06296 | 0.00226 |
| Concordance_oral_16S | unifrac_weighted_phyloseq | 0.86975 | 0.0435 | 0.00283 |
| covid | unifrac_weighted_phyloseq | 1.85404 | 0.19973 | 0.00882 |
| HIV_gut | unifrac_weighted_phyloseq | 0.93881 | 0.07099 | 0.00786 |
| IBD_16s | unifrac_weighted_phyloseq | 0.98643 | 0.07596 | 0.01034 |
| IBD_wgs | unifrac_weighted_phyloseq | 1.02117 | 0.08402 | 0.00728 |
| Pig_gut | unifrac_weighted_phyloseq | 10.92424 | 0.20072 | 0.18948 |

**Table 8** Results without applying any remedial measures for each dissimilarity measure across different datasets. Each row shows the PERMANOVA pseudo-$F$ statistic, pseudo-$R^2$, and MiRKAT $R^2$ values computed without Higham correction or Tikhonov regularization.

| | Dissimilarity | pseudo_F | pseudo_R2 | MiRKAT_Rsquared |
|---|---|---|---|---|
| cardiovascular | bray | 1.74914 | 0.07513 | 0.0035 |
| Concordance_oral_16S | bray | 1.01089 | 0.0502 | 0.00565 |
| covid | bray | 1.45796 | 0.16406 | 0.01458 |
| HIV_gut | bray | 1.26877 | 0.09361 | 0.01496 |
| IBD_16s | bray | 1.45303 | 0.10801 | 0.0139 |
| IBD_wgs | bray | 1.36777 | 0.10941 | 0.01366 |
| Pig_gut | bray | 5.46714 | 0.11165 | 0.09125 |
| cardiovascular | cao | 2.03228 | 0.08624 | 0.00414 |
| Concordance_oral_16S | cao | 1.1542 | 0.05692 | 0.00581 |
| HIV_gut | cao | 1.18867 | 0.08822 | 0.01569 |
| IBD_16s | cao | 1.36084 | 0.10185 | 0.00931 |
| IBD_wgs | cao | 1.27353 | 0.10265 | 0.01426 |
| cardiovascular | chao | 1.25552 | 0.05509 | 0.00308 |
| Concordance_oral_16S | chao | 1.15665 | 0.05703 | 0.00763 |
| covid | chao | 1.76348 | 0.19185 | 0.01377 |
| HIV_gut | chao | 1.29223 | 0.09517 | 0.01629 |
| IBD_16s | chao | 1.47969 | 0.10977 | 0.01469 |
| IBD_wgs | chao | 1.30062 | 0.1046 | 0.0199 |
| Pig_gut | chao | 5.59982 | 0.11405 | 0.09188 |
| cardiovascular | jaccard | 1.51569 | 0.06576 | 0.0033 |
| Concordance_oral_16S | jaccard | 1.00831 | 0.05008 | 0.00597 |
| Concordance_oral_16S | unifrac_unweighted | 1.24972 | 0.06134 | 0.00678 |
| cardiovascular | unifrac_weighted_gunifrac | 2.04262 | 0.08664 | 0.00458 |
| Concordance_oral_16S | unifrac_weighted_gunifrac | 1.00144 | 0.04976 | 0.00416 |
| covid | unifrac_weighted_gunifrac | 1.27994 | 0.14698 | 0.01511 |
| HIV_gut | unifrac_weighted_gunifrac | 1.12547 | 0.08392 | 0.01154 |
| IBD_16s | unifrac_weighted_gunifrac | 1.65212 | 0.12102 | 0.02231 |
| IBD_wgs | unifrac_weighted_gunifrac | 1.47011 | 0.11664 | 0.02729 |
| Pig_gut | unifrac_weighted_gunifrac | 9.42701 | 0.17811 | 0.15373 |
| cardiovascular | unifrac_weighted_phyloseq | 1.67729 | 0.07226 | 0.00305 |
| Concordance_oral_16S | unifrac_weighted_phyloseq | 1.00144 | 0.04976 | 0.00416 |
| covid | unifrac_weighted_phyloseq | 2.01959 | 0.21375 | 0.01178 |
| HIV_gut | unifrac_weighted_phyloseq | 1.02521 | 0.07702 | 0.009 |
| IBD_16s | unifrac_weighted_phyloseq | 1.11338 | 0.0849 | 0.01202 |
| IBD_wgs | unifrac_weighted_phyloseq | 1.12748 | 0.09196 | 0.0115 |
| Pig_gut | unifrac_weighted_phyloseq | 11.67488 | 0.2116 | 0.20037 |

**Table 9** Results after applying Higham correction to each dissimilarity measure across different datasets. Each row shows the PERMANOVA pseudo-$F$ statistic, pseudo-$R^2$, and MiRKAT $R^2$ values following Higham matrix adjustment.

| | Dissimilarity | pseudo_F | pseudo_R2 | MiRKAT_Rsquared |
|---|---|---|---|---|
| cardiovascular | bray | 1.44935 | 0.06306 | 0.00324 |
| Concordance_oral_16S | bray | 1.07054 | 0.05301 | 0.00585 |
| covid | bray | 1.43635 | 0.16203 | 0.01475 |
| HIV_gut | bray | 1.24717 | 0.09216 | 0.01423 |
| IBD_16s | bray | 1.32115 | 0.09918 | 0.011 |
| IBD_wgs | bray | 1.27547 | 0.10279 | 0.01112 |
| Pig_gut | bray | 5.39175 | 0.11028 | 0.08984 |
| cardiovascular | cao | 1.33273 | 0.05828 | 0.00329 |
| Concordance_oral_16S | cao | 1.1355 | 0.05604 | 0.00595 |
| HIV_gut | cao | 1.16712 | 0.08676 | 0.01296 |
| IBD_16s | cao | 1.12454 | 0.08568 | 0.00616 |
| IBD_wgs | cao | 1.22063 | 0.0988 | 0.01202 |
| cardiovascular | chao | 1.07329 | 0.04748 | 0.00297 |
| Concordance_oral_16S | chao | 1.13508 | 0.05603 | 0.00631 |
| covid | chao | 1.37005 | 0.15571 | 0.01634 |
| HIV_gut | chao | 1.18032 | 0.08765 | 0.01193 |
| IBD_16s | chao | 1.1305 | 0.0861 | 0.00679 |
| IBD_wgs | chao | 1.07249 | 0.08787 | 0.00646 |
| Pig_gut | chao | 2.69382 | 0.05832 | 0.03566 |
| cardiovascular | jaccard | 1.51477 | 0.06572 | 0.0033 |
| Concordance_oral_16S | jaccard | 1.01575 | 0.05043 | 0.00599 |
| Concordance_oral_16S | unifrac_unweighted | 1.24003 | 0.06089 | 0.00673 |
| cardiovascular | unifrac_weighted_gunifrac | 1.26428 | 0.05546 | 0.00328 |
| Concordance_oral_16S | unifrac_weighted_gunifrac | 1.09925 | 0.05435 | 0.00569 |
| covid | unifrac_weighted_gunifrac | 1.25596 | 0.14462 | 0.0156 |
| HIV_gut | unifrac_weighted_gunifrac | 1.14088 | 0.08497 | 0.01105 |
| IBD_16s | unifrac_weighted_gunifrac | 1.30348 | 0.09798 | 0.01254 |
| IBD_wgs | unifrac_weighted_gunifrac | 1.18906 | 0.0965 | 0.01227 |
| Pig_gut | unifrac_weighted_gunifrac | 6.45844 | 0.12928 | 0.1051 |
| cardiovascular | unifrac_weighted_phyloseq | 1.2756 | 0.05593 | 0.00299 |
| Concordance_oral_16S | unifrac_weighted_phyloseq | 1.09925 | 0.05435 | 0.00569 |
| covid | unifrac_weighted_phyloseq | 1.39239 | 0.15785 | 0.01495 |
| HIV_gut | unifrac_weighted_phyloseq | 1.12523 | 0.0839 | 0.01035 |
| IBD_16s | unifrac_weighted_phyloseq | 1.0691 | 0.0818 | 0.00598 |
| IBD_wgs | unifrac_weighted_phyloseq | 1.07024 | 0.0877 | 0.00577 |
| Pig_gut | unifrac_weighted_phyloseq | 4.92544 | 0.10171 | 0.08364 |

**Table 10** Results after applying Tikhonov regularization to each dissimilarity measure across different datasets. Each row shows the PERMANOVA pseudo-$F$ statistic, pseudo-$R^2$, and MiRKAT $R^2$ values following Tikhonov adjustment.

### References


[1] Whittaker, R.H.: Vegetation of the siskiyou mountains, oregon and california. Ecological monographs **30**(3), 279–338 (1960)

[2] Pane, C., Sorrentino, R., Scotti, R., Molisso, M., Di Matteo, A., Celano, G., Zaccardelli, M.: Alpha and beta-diversity of microbial communities associated to plant disease suppressive functions of on-farm green composts. Agriculture **10**(4), 113 (2020)

[3] Koleff, P., Gaston, K.J., Lennon, J.J.: Measuring beta diversity for presence–absence data. Journal of Animal Ecology **72**(3), 367–382 (2003)

[4] Socolar, J.B., Gilroy, J.J., Kunin, W.E., Edwards, D.P.: How should beta-diversity inform biodiversity conservation? Trends in ecology & evolution **31**(1), 67–80 (2016)

[5] Bush, A., Harwood, T., Hoskins, A.J., Mokany, K., Ferrier, S.: Current uses of beta-diversity in biodiversity conservation: A response to. Trends Ecol Evol **31**, 337 (2016)

[6] De Cáceres, M., Legendre, P., Valencia, R., Cao, M., Chang, L.-W., Chuyong, G., Condit, R., Hao, Z., Hsieh, C.-F., Hubbell, S., *et al.*: The variation of tree beta diversity across a global network of forest plots. Global Ecology and Biogeography **21**(12), 1191–1202 (2012)

[7] Prober, S.M., Leff, J.W., Bates, S.T., Borer, E.T., Firn, J., Harpole, W.S., Lind, E.M., Seabloom, E.W., Adler, P.B., Bakker, J.D., *et al.*: Plant diversity predicts beta but not alpha diversity of soil microbes across grasslands worldwide. Ecology letters **18**(1), 85–95 (2015)

[8] Villarino, E., Watson, J.R., Chust, G., Woodill, A.J., Klempay, B., Jonsson, B., Gasol, J.M., Logares, R., Massana, R., Giner, C.R., *et al.*: Global beta diversity patterns of microbial communities in the surface and deep ocean. Global Ecology and Biogeography **31**(11), 2323–2336 (2022)

[9] Jaccard, P.: Étude comparative de la distribution florale dans une portion des alpes et des jura. Bull Soc Vaudoise Sci Nat **37**, 547–579 (1901)

[10] Kosub, S.: A note on the triangle inequality for the jaccard distance. Pattern Recognition Letters **120**, 36–38 (2019)

[11] Lipkus, A.H.: A proof of the triangle inequality for the tanimoto distance. Journal of Mathematical Chemistry **26**(1), 263–265 (1999)

[12] Levandowsky, M., Winter, D.: Distance between sets. Nature **234**(5323), 34–35 (1971)

[13] Ioffe, S.: Improved consistent sampling, weighted minhash and l1 sketching. In: 2010 IEEE International Conference on Data Mining, pp. 246–255 (2010). IEEE

[14] Taha, A.A., Hanbury, A.: Metrics for evaluating 3d medical image segmentation: analysis, selection, and tool. BMC medical imaging **15**, 1–28 (2015)

[15] Kulczyński, S.: Die Pflanzenassoziationen der Pieninen vol. 3. Imprimerie de l'Université, Lwów (1928)

[16] Legendre, P.: Numerical Ecology. Elsevier, Amsterdam (2012)

[17] Deza, M.-M., Deza, E.: Dictionary of Distances. Elsevier, Amsterdam (2006)

[18] Shi, G.: A comparative study of 39 binary similarity coefficients (1993)

[19] Cheetham, A.H., Hazel, J.E.: Binary (presence-absence) similarity coefficients. Journal of Paleontology, 1130–1136 (1969)

[20] Hausdorf, B., Hennig, C.: Biotic element analysis in biogeography. Systematic Biology **52**(5), 717–723 (2003)

[21] Hennig, C., Hausdorf, B.: Distance-based parametric bootstrap tests for clustering of species ranges. Computational Statistics & Data Analysis **45**(4), 875–895 (2004)

[22] Hennig, C., Hausdorf, B.: A robust distance coefficient between distribution areas incorporating geographic distances. Systematic biology **55**(1), 170–175 (2006)

[23] Bray, J.R., Curtis, J.T.: An ordination of the upland forest communities of southern wisconsin. Ecological monographs **27**(4), 326–349 (1957)

[24] Dice, L.R.: Measures of the amount of ecologic association between species. Ecology **26**(3), 297–302 (1945)

[25] Sorensen, T.: A method of establishing groups of equal amplitude in plant sociology based on similarity of species content and its application to analyses of the vegetation on danish commons. Biologiske skrifter **5**, 1–34 (1948)

[26] Greenacre, M., Primicerio, R.: Chapter: Measures of distance between samples: Non-euclidean. Universitat Pompeu Fabra Barcelona Department of Economics and Business [Online]. http://www. econ. upf. edu/-michael/stanford/maeb5. pdf (2008)

[27] Greenacre, M.: 'size'and 'shape'in the measurement of multivariate proximity. Methods in Ecology and Evolution **8**(11), 1415–1424 (2017)

[28] Lozupone, C., Lladser, M.E., Knights, D., Stombaugh, J., Knight, R.: Unifrac: an effective distance metric for microbial community comparison. The ISME journal

**5**(2), 169–172 (2011)

[29] Lozupone, C., Knight, R.: Unifrac: a new phylogenetic method for comparing microbial communities. Applied and environmental microbiology **71**(12), 8228–8235 (2005)

[30] Xia, Y., Sun, J.: Beta diversity metrics and ordination. In: Bioinformatic and Statistical Analysis of Microbiome Data: From Raw Sequences to Advanced Modeling with QIIME 2 and R, pp. 335–395. Springer, New York (2023)

[31] Chen, J., Bittinger, K., Charlson, E.S., Hoffmann, C., Lewis, J., Wu, G.D., Collman, R.G., Bushman, F.D., Li, H.: Associating microbiome composition with environmental covariates using generalized unifrac distances. Bioinformatics **28**(16), 2106–2113 (2012)

[32] Lozupone, C.A., Hamady, M., Kelley, S.T., Knight, R.: Quantitative and qualitative $\beta$ diversity measures lead to different insights into factors that structure microbial communities. Applied and environmental microbiology **73**(5), 1576–1585 (2007)

[33] Wong, R.G., Wu, J.R., Gloor, G.B.: Expanding the unifrac toolbox. PloS one **11**(9), 0161196 (2016)

[34] Minkowski, H.: Geometrie der Zahlen vol. 1. BG Teubner, Leipzig (1910)

[35] Şuhubi, E.S.: Metric Spaces, pp. 261–356. Springer, Dordrecht (2003). https://doi.org/10.1007/978-94-017-0141-9_5

[36] Heath, T.L.: The Thirteen Books of Euclid's Elements. Dover Publications, Inc, New York (1956)

[37] Elmore, K.L., Richman, M.B.: Euclidean distance as a similarity metric for principal component analysis. Monthly weather review **129**(3), 540–549 (2001)

[38] Korenius, T., Laurikkala, J., Juhola, M.: On principal component analysis, cosine and euclidean measures in information retrieval. Information Sciences **177**(22), 4893–4905 (2007)

[39] Greenacre, M., Groenen, P.J., Hastie, T., d'Enza, A.I., Markos, A., Tuzhilina, E.: Principal component analysis. Nature Reviews Methods Primers **2**(1), 100 (2022)

[40] Singh, A., Yadav, A., Rana, A.: K-means with three different distance metrics. International Journal of Computer Applications **67**(10) (2013)

[41] Kapil, S., Chawla, M.: Performance evaluation of k-means clustering algorithm with various distance metrics. In: 2016 IEEE 1st International Conference on Power Electronics, Intelligent Control and Energy Systems (ICPEICES), pp. 1–4 (2016). IEEE

[42] Xu, J., Chi, E., Lange, K.: Generalized linear model regression under distance-to-set penalties. Advances in neural information processing systems **30** (2017)

[43] Krislock, N., Wolkowicz, H.: Euclidean Distance Matrices and Applications. Springer, New York (2012)

[44] Bishop, C.M.: Pattern recognition and machine learning. Springer google schola **2**, 1122–1128 (2006)

[45] Deza, E., Deza, M.M., Deza, M.M., Deza, E.: Encyclopedia of Distances. Springer, New York (2009)

[46] Chaya, C., Castura, J., Greenacre, M.: One citation, one vote! a new approach for analysing check-all-that-apply (cata) data, using l1 norm methods. arXiv preprint arXiv:2502.15945 (2025)

[47] Jiang, X., Hu, X., He, T.: Identification of the clustering structure in microbiome data by density clustering on the manhattan distance. Science China Information Sciences **59**, 1–7 (2016)

[48] Pearson, K.: X. on the criterion that a given system of deviations from the probable in the case of a correlated system of variables is such that it can be reasonably supposed to have arisen from random sampling. The London, Edinburgh, and Dublin Philosophical Magazine and Journal of Science **50**(302), 157–175 (1900)

[49] Dodge, Y.: The Concise Encyclopedia of Statistics. Springer, New York (2008)

[50] Liu, X., Wang, D.: A spectral histogram model for texton modeling and texture discrimination. Vision Research **42**(23), 2617–2634 (2002)

[51] Dixon, P.: Vegan, a package of r functions for community ecology. Journal of vegetation science **14**(6), 927–930 (2003)

[52] Hellinger, E.: Neue begründung der theorie quadratischer formen von unendlichvielen veränderlichen. Journal für die reine und angewandte Mathematik **1909**(136), 210–271 (1909)

[53] Zhu, C., Xiao, F.: A belief hellinger distance for d–s evidence theory and its application in pattern recognition. Engineering Applications of Artificial Intelligence **106**, 104452 (2021)

[54] Vázquez-Castellanos, J.F.: Diversity analysis in viral metagenomes. The Human Virome: Methods and Protocols, 203–230 (2018)

[55] Markatou, M., Karlis, D., Ding, Y.: Distance-based statistical inference. Annual Review of Statistics and Its Application **8**(1), 301–327 (2021)

[56] Karagrigoriou, A.: Statistical inference: The minimum distance approach. Taylor

& Francis (2012)

[57] Cieslak, D.A., Hoens, T.R., Chawla, N.V., Kegelmeyer, W.P.: Hellinger distance decision trees are robust and skew-insensitive. Data Mining and Knowledge Discovery **24**, 136–158 (2012)

[58] Torgersen, E.: Comparison of Statistical Experiments vol. 36. Cambridge University Press, Cambridge, UK (1991)

[59] Mahalanobis, P.C.: On the generalized distance in statistics. Sankhyā: The Indian Journal of Statistics, Series A (2008-) **80**, 1–7 (2018)

[60] Manly, B.F., Manly, B.: Multivariate statistical methods (1994)

[61] Oksanen, J., Kindt, R., Legendre, P., O'Hara, B., Stevens, M.H.H., Oksanen, M.J., Suggests, M.: The vegan package. Community ecology package **10**(631-637), 719 (2007)

[62] McMurdie, P.J., Holmes, S.: phyloseq: an r package for reproducible interactive analysis and graphics of microbiome census data. PloS one **8**(4), 61217 (2013)

[63] Pavoine, S., Dufour, A.-B., Chessel, D.: From dissimilarities among species to dissimilarities among communities: a double principal coordinate analysis. Journal of theoretical biology **228**(4), 523–537 (2004)

[64] Ramette, A.: Multivariate analyses in microbial ecology. FEMS microbiology ecology **62**(2), 142–160 (2007)

[65] Aitchison, J.: The statistical analysis of compositional data. Journal of the Royal Statistical Society: Series B (Methodological) **44**(2), 139–160 (1982)

[66] Pawlowsky-Glahn, V., Egozcue, J.J., Tolosana-Delgado, R.: Modeling and Analysis of Compositional Data. John Wiley & Sons, Chichester, UK (2015)

[67] Zhang, Y., Schluter, J., Zhang, L., Cao, X., Jenq, R.R., Feng, H., Haines, J., Zhang, L.: Review and revamp of compositional data transformation: A new framework combining proportion conversion and contrast transformation. Computational and Structural Biotechnology Journal **23**, 4088–4107 (2024)

[68] Martino, C., Morton, J.T., Marotz, C.A., Thompson, L.R., Tripathi, A., Knight, R., Zengler, K.: A novel sparse compositional technique reveals microbial perturbations. MSystems **4**(1), 10–1128 (2019)

[69] Resnick, S.I.: A Probability Path. Modern Probability Theory. Birkhäuser, Boston, MA (1999). https://doi.org/10.1007/978-1-4612-0537-1
